# Habitat loss does not always entail negative genetic consequences

**DOI:** 10.1101/528430

**Authors:** Carolina S. Carvalho, Éder C. M. Lanes, Amanda R. Silva, Cecilio F. Caldeira, Nelson Carvalho-Filho, Markus Gastauer, Vera L. Imperatriz-Fonseca, Wilson Nascimento Júnior, Guilherme Oliveira, José O. Siqueira, Pedro L. Viana, Rodolfo Jaffé

## Abstract

Although habitat loss has large, consistently negative effects on biodiversity, its genetic consequences are not yet fully understood. This is because measuring the genetic consequences of habitat loss requires accounting for major methodological limitations like the confounding effect of habitat fragmentation, historical processes underpinning genetic differentiation, time-lags between the onset of disturbances and genetic outcomes, and the need for large numbers of samples, genetic markers and replicated landscapes to ensure sufficient statistical power. In this paper we overcame all these challenges to assess the genetic consequences of extreme habitat loss driven by mining in two herbs endemic to Amazonian Savannas. Relying on genotyping-by-sequencing of hundreds of individuals collected across two mining landscapes, we identified thousands of neutral and independent single-nucleotide polymorphisms (SNPs) in each species and used these to evaluate population structure, genetic diversity and gene flow. Since open-pit mining in our study region rarely involves habitat fragmentation, we were able to assess the independent effect of habitat loss. We also accounted for the underlying population structure when assessing landscape effects on genetic diversity and gene flow, assessed the sensitivity of our analyses to the resolution of spatial data, and used annual species and cross-year analyses to minimize and quantify possible time-lag effects. We found that both species are remarkably resilient, as genetic diversity and gene flow patterns were unaffected by habitat loss. Whereas historical habitat amount was found to influence inbreeding; heterozygosity and inbreeding were not affected by habitat loss in either species, and gene flow was mainly influenced by geographic distance, pre-mining land cover and local climate. Our study demonstrates that it is not possible to generalize about the genetic consequences of habitat loss, and imply that future conservation efforts need to consider species-specific genetic information.

## Introduction

In spite of ample evidence showing that habitat loss has large, consistently negative effects on biodiversity (Fahrig, 2003), very few studies have assessed the consequences of habitat amount on genetic variation (DiLeo and Wagner, 2016; Monteiro et al., 2019). Habitat loss can potentially impact the demographics of natural populations, reducing population size, gene flow and genetic diversity, and thereby increasing inbreeding and extinction risk (Allendorf et al., 2013). Understanding the genetic consequences of habitat loss is therefore essential to safeguard biological diversity and fulfill Aichi Biodiversity Targets and Sustainable Development Goals (Tittensor et al., 2014).

Important limitations constrain the quantification of habitat amount effects on genetic variation. Firstly, habitat loss and fragmentation are often confounded, so disentangling the relative contribution of habitat amount requires controlling for fragmentation (Fahrig, 2003). Secondly, landscape effects can also be easily confounded with historical demographic processes that underlying population structure (Llorens et al., 2018). Thirdly, a coarse resolution of spatial data and time-lags between the onset of disturbances and genetic responses may mask the effects of recent landscape modification (Anderson et al., 2010; Balkenhol et al., 2016). Finally, large numbers of samples and genetic markers, and replicated sampling designs that capture enough landscape heterogeneity are needed to detect or rule out possible landscape effects with sufficient statistical power (McCartney-Melstad et al., 2018; Storfer et al., 2010). Failure in overcoming any of these limitations may hide important detrimental effects to the maintenance of genetic variability, or reveal spurious patterns unrelated to habitat loss.

Few studies have attempted to quantify the impact of habitat loss on both genetic diversity and gene flow, and neither has yet accounted for all the methodological limitations outlined above (Balkenhol et al., 2016; DiLeo and Wagner, 2016). Here we fill this important knowledge gap assessing the genetic consequences of extreme habitat loss driven by open-pit mining in two endemic plants from the Eastern Amazon. Firstly, we were able to assess the independent effect of habitat loss, as open-pit mining in our study region rarely involves habitat fragmentation (See Fig. S1 in Supporting Information). We also controlled for historical demographic processes by accounting for the underlying population structure when assessing landscape effects on genetic diversity and gene flow; assessed the sensitivity of our analyses to the resolution of spatial data; and used annual species (which complete a full reproductive cycle and die within one year) to minimize possible time-lag effects. Finally, we sampled hundreds of individuals scattered across two separate regions exposed to mining, and genotyped them at thousands of single nucleotide polymorphisms (SNPs) distributed across their genomes to assure high statistical power.

The banded iron formations known as Cangas from the Carajás Mineral Province in the Eastern Amazon harbor one of the world’s largest deposits of high-grade iron ore (Skirycz et al., 2014), which has attracted substantial attention from mining companies. In fact two of the world’s largest iron-ore mines are located in the region (Fig. 1), with operations in Serra Norte dating back to the 1980s, while Serra Sul only began activities in 2014. Based on a curated inventory of Canga plants from this region (Viana et al., 2016), we selected annual herbs (to minimize time-lag effects), occurring exclusively in Canga ecosystems (where mining activities are concentrated), and endemic to our study region. From the few available species meeting these criteria, *Brasilianthus carajensis* (Melastomataceae) and *Monogereion carajensis* (Asteraceae), were among the easiest to find and identify in the field. Since both species seem to be pollinated by insects and their seeds dispersed by the wind (Cruz et al., 2016; Rocha et al., 2017), we expected them to be susceptible to habitat loss, given that the progeny of insect- and wind-pollinated plants has been shown to be strongly negatively affected by habitat fragmentation (Aguilar et al., 2019). Relying on genotyping-by-sequencing of hundreds of individuals from both species collected across these two mining landscapes, we assessed the influence of habitat loss on genetic diversity and gene flow. We also performed a suit of germination experiments to determine if these plants are able to successfully colonize iron-ore mines. We predicted that: i) Individuals surrounded by undisturbed habitats would show higher genetic diversity and lower inbreeding than those exposed to habitat loss driven by mining; ii) Gene flow would be best explained by recent landscape modifications, and mining areas would represent barriers to gene flow; iii) Mining waste substrates would hinder germination.

**Fig. 1.**
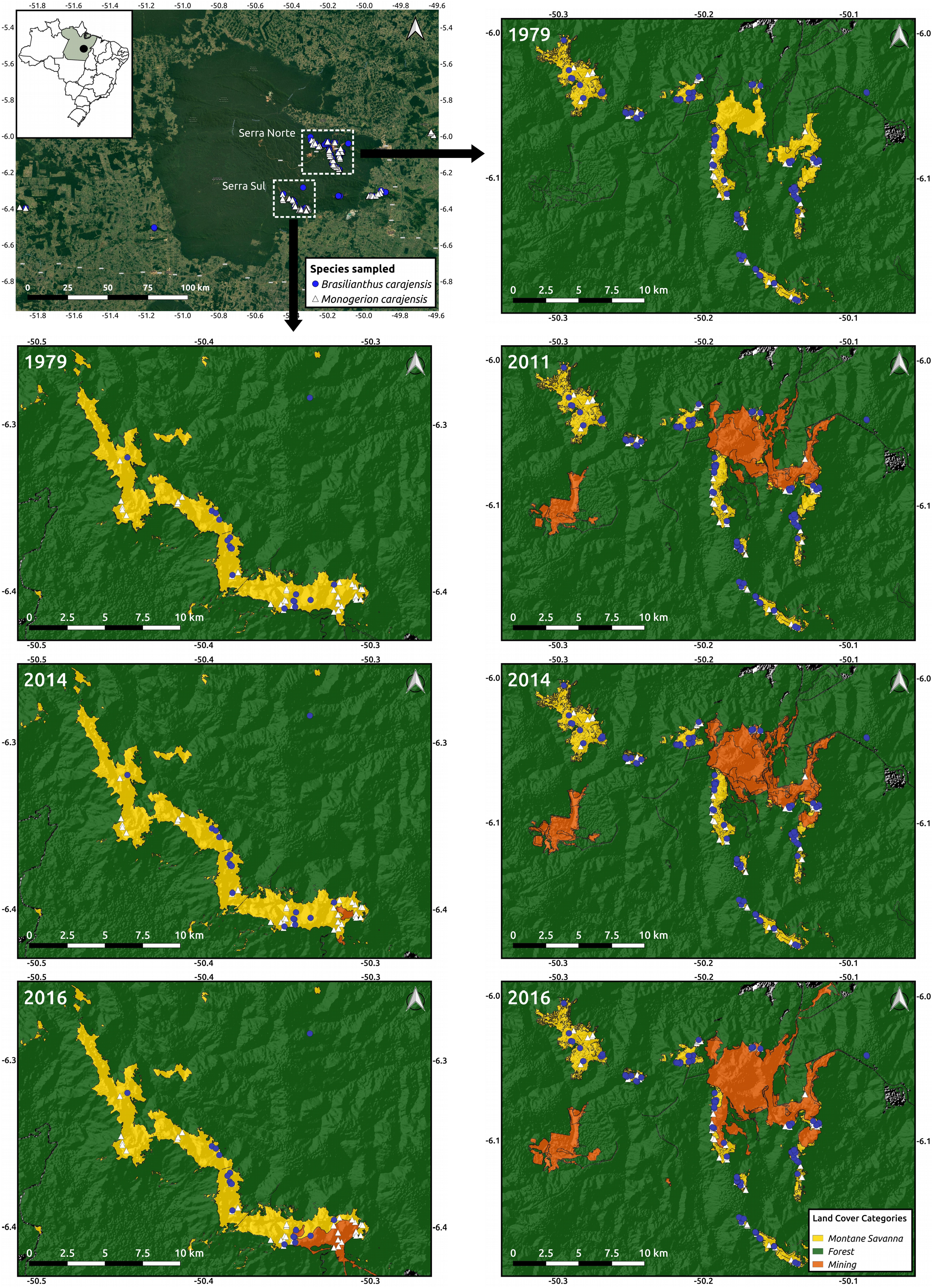
Map of the study region depicting the location of the collected samples from *Brasilianthus carajensis* (blue circles) and *Monogereion carajensis* (white triangles) in Serra Norte (right panels) and Serra Sul (left panels). Hill shade maps are shown overlaid with land cover color maps for the different years analyzed. The location of the Carajás Mineral Province within Brazil is shown on the upper left corner.

## Materials and Methods

### Sampling, DNA extraction, and genome size estimation

We collected leaf tissue samples of 150 individuals of *B. carajensis* and 207 individuals of *M. carajensis* between February and June 2017 (SISBIO collection permit N. 48272-4). Samples were collected in the main Canga plateaus of our study area, comprising the entire occurrence range of both species. Care was taken to sample individuals at or around iron ore mines and separated by at least 20 m from each other to avoid collecting siblings (Fig. 1). Both species exhibit a patchy distribution, with *B. carajensis* usually occurring in aggregates of up to hundreds of individuals occupying rocky exposed soils, while isolated individuals of *M. carajensis* were often found in shaded areas and soils with a larger organic mater content (personal observation). Our study area harbors some of the world’s largest iron-ore mines, one located in Serra Norte, comprised of an archipelago of Canga plateaus, and one in Serra Sul formed by a large continuous plateau. While in Serra Norte we collected 103 and 120 individuals, in Serra Sul we sampled 17 and 149 individuals of *B. carajensis* and *M. carajensis* respectively. To ensure high DNA quality and concentration, we preserved *B. carajensis* samples in silica and *M. carajensis* samples in 10 mL of a NaCl-saturated solution of 2% CTAB (Rogstad, 1992), and stored them at −80 °C until analysis. Total DNA of *B. carajensis* was extracted using a CTAB 2% protocol (Doyle and Doyle, 1987) followed by a DNA purification protocol (Michaels et al., 1994); whereas the DNeasy Plant Mini Kit (Qiagen, EUA) was used for *M. carajensis.* DNA concentration for both species was quantified using the Qubit High SensitivityAssay kit (Invitrogen), and DNA integrity assessed through 1.2% agarose gel electrophoresis. All DNA samples were adjusted to a final concentration of 5 ng/μL in a final volume of 30 μL. We used flow cytometry to estimate genome size in both species. Nuclei were obtained from fresh leaf tissues chopped along with references in general purpose buffer with 1% Triton X-100 and 1% PVP-30 (Loureiro et al., 2007). The whole sample preparation was conducted on ice until the events acquisition on a PI fluorescence mean under a 575/26 bandpass filter. Triplicates of 1000 PI stained nuclei were analyzed under a 488 nm laser on BD FACS Aria II cytometer. The internal standard used was tomato (*Lycopersicon esculentum*; 2C = 1.98 pg, (Dolezel et al., 1992)).

### RAD sequencing and SNP discovery

DNA samples were shipped to SNPSaurus (http://snpsaurus.com/) for sequencing and bioinformatic analyses of raw reads (trimming and variant calling). Briefly, nextRAD genotyping-by-sequencing libraries were prepared (Russello et al., 2015) using Nextera DNA Flex reagent (Illumina, Inc) and considering the estimated genome size of each species (2C DNA content was 508 Mbp in *B. carajensis* and 6,284 Mbp in *M. carajensis*). Fragmented DNA was then amplified for 25 cycles at 75 degrees, with one of the primers matching the adapter and extending 8 nucleotides into the genomic DNA with the selective sequences GTGTAGAA (*B. carajensis*) and TGCAGGAG (*M. carajensis*). Thus, only fragments starting with a sequence that can be hybridized by the selective sequence of the primer will be efficiently amplified. The nextRAD libraries were sequenced on a HiSeq 4000 with four lanes of 150 bp reads (University of Oregon). Genotyping analysis used custom scripts (SNPsaurus, LLC) that trimmed the reads using bbduk (BBMap tools, http://sourceforge.net/projects/bbmap/): bbmap/bbduk.sh in=$file out=$outfile ktrim=r k=17 hdist=1 mink=8 ref=bbmap/resources/ nextera.fa.gz minlen=100 ow=t qtrim=r trimq=10. A de novo reference was created by collecting 10 million reads in total, evenly from the samples, and excluding reads that had counts fewer than 5 or more than 700 for *B. carajensis*; and fewer than 6 or more than 1,000 for *M. carajensis*. The remaining loci were then aligned to each other to identify allelic loci and collapse allelic haplotypes to a single representative. All reads were mapped to the reference with an alignment identity threshold of 90% using bbmap (BBMap tools). Genotype calling was done using Samtools and bcftools (samtools mpileup -gu -Q 12 -t DP, DPR -f ref.fasta -b samples.txt | bcftools call -cv - > genotypes.vcf). The vcf was filtered to remove alleles with a population frequency of less than 3%. Loci were removed that were heterozygous in all samples or had more than 2 alleles in a sample (suggesting collapsed paralogs). The absence of artifacts was checked by counting SNPs at each read nucleotide position and determining that SNP number did not increase with reduced base quality at the end of the read. A total of 43,887 contigs were generated for *B. carajensis* (sequencing depth ranged between 18 and 239), and 36,040 for *M. carajensis* (depth ranging between 14 and 246). Geographic coordinates in decimal degrees, genotypes in Variant Call Format and sequences in FASTA format for both species are provided here: https://figshare.com/articles/Habitat_loss_does_not_always_entail_negative_genetic_consequences/8224175.

### Neutral datasets

The R package *r2vcftools* (https://github.com/nspope/r2vcftools) - a wrapper for VCFtools (Danecek et al., 2011) - was used to perform final quality control on the genotype data (see detailed script here: https://github.com/rojaff/r2vcftools_basics. We excluded loci containing more than 30% missing data, and filtered them for quality (Phred score > 50 both species), read depth (30 – 240 both species), linkage disequilibrium (LD, *r*^2^ < 0.6 and *r*^2^ < 0.4 for *B. carajensis* and *M. carajensis*, respectively), and strong deviations from the Hardy Weinberg Equilibrium (HWE, *p* < 0.0001 both species) (O’Leary et al., 2018). Additionally, we removed any potential loci under selection detected through genome scans, whereby *F*_*ST*_ outlier tests were applied after assessing population structure with the *snmf* function from the LEA package (Frichot et al., 2014). The genomic inflation factor and a trial-and-error approach were used to calibrate *p*-values, and the Benjamini-Hochberg algorithm (*q* = 0.05) was used to correct for false discovery rates (François et al., 2016), see example script here: http://membres-timc.imag.fr/Olivier.Francois/LEA/files/LEA_snmf.html). The resulting sets of neutral and independent loci were then used in all subsequent analyses.

### Genetic structure

We used two complementary genetic clustering method to assess population structure: Admixture *(Alexander et al., 2009)* and Discriminant Analysis of Principal Components - DAPC from the *adegenet* package (Jombart et al., 2010; Jombart and Ahmed, 2011). Admixture puts individuals into groups to maximize HWE while that DAPC minimizes allelic differences. For the former analysis, the number of ancestral populations (*k*) was allowed to vary between 1 and 10, and the best *k* was chosen based on cross-validation errors (Frichot et al., 2014). For the second analyses, the number of clusters was assessed using the function “*find.cluster*”, which runs successive *k*-means clustering with an increasing number of clusters, and then determined the best-supported number of genetic clusters using the Bayesian Information Criterion (BIC). Considering the ancestry coefficients assigned by Admixture, we then estimated expected heterozygosity (*H*_*E*_) and inbreeding coefficients (*F*) for each genetic cluster using the “het” option in VCFtools implemented in *r2vcftools* (Danecek et al., 2011). Additionally, we assessed fine-scale spatial genetic structure within each genetic cluster by quantifying spatial autocorrelation in Yang’s genetic relatedness between pairs of individuals *(Yang et al., 2010)*. To do so we used local polynomial fitting (LOESS) of pairwise relatedness and pairwise geographic distance. In order to test if the average observed relatedness predicted by LOESS at a given distance differs from the null model, row and column indices for the relatedness matrix were permuted 999 times. The smoothing parameter was fixed as the default for loess() in R (span = 0.75). At each permutation a LOESS model was re-fitted using the permuted relatedness and geographic distance matrix, and 95 % percentiles of the permutation-derived LOESS predictions were used to generate confidence envelopes around the null expectation (see example script here: https://github.com/rojaff/Lplot).

### Land cover maps

To account for time-lag effects when assessing the genetic consequences of habitat loss, we built land cover maps for different years (2016, 2014, 2011 and 1979), comprising pre-mining maps (1979). *Landsat* 2 images (spatial resolution of 80 meters in 7 spectral bands) were used for year 1979, *Landsat* 7 images (spatial resolution of 30 meters in 7 spectral bands) for year 2011, and *Sentinel* images (spatial resolution of 10 meters in 4 spectral bands) for years 2014 and 2016. Images were downloaded from the Earth Explorer Server (https://earthexplorer.usgs.gov/), selecting scenes from the month of July to minimize clouds. All images were converted to ground reflectance in percentage using the ATCOR algorithm of the PCI Geomatica 2016 software. The scenes were joined to create a mosaic of the study area and derive the Normalized Difference Vegetation Index - NDVI (Tarpley et al., 1984). We then employed the eCognition 9 software using a Geographic Object-Based Image Analysis (GEOBIA) to classify land cover types. The Multi-resolution classification algorithm was selected, given that it allows obtaining segments with different sizes due to brightness, shape, smoothness and compactness. Montane Savanna (Canga), Water, Forest, Mine, Pasture and Urban classes were identified.

### Genetic diversity

To assess the effect of habitat loss on genetic diversity, we regressed individual-level diversity metrics (*H* and *f*), estimated with “het” option in VCFtools (Danecek et al., 2011), on historical habitat amount and habitat loss (measured in area) driven by mining in different years, using high resolution land cover maps (10 × 10m). By so doing we explicitly evaluated the effect of habitat loss within a buffer accounting for historical habitat amount. Historical habitat amount was calculated by extracting the proportion of Canga habitat in a buffer surrounding each individual using pre-mining maps (1979). Habitat loss in different years (2011, 2014, and 2016) was calculated by subtracting habitat amount for a given year from historical habitat amount. To select an optimal buffer size we first ran uni-variate models using habitat amount extracted from the most recent land cover maps (2016) with buffers varying in size between 100 and 900m, and then compared all models using AIC. As habitat amount calculated with the largest buffers (900m) was always among the best models (ΔAIC ≤ 2), we chose this buffer size to encompass a greater portion of lost areas (Table S1). Additionally, the small flowers of our study species suggest their pollinators do not forage beyond 1Km (Greenleaf et al., 2007). In Serra Norte, 44% of individuals of *B. carajensis* and 41% of *M. carajensis* experienced habitat loss, whereas 55% and 57% of individuals of *B. carajensis* and *M. carajensis*, respectively, faced habitat loss in Serra Sul.

Because genetic parameters (*H* and *f*) are affected by demographic processes, we explicitly accounted for demography either by analyzing individuals belonging to the same cluster or by including genetic cluster identity as a random effect in our models. In Serra Norte, which comprises an archipelago of Canga plateaus, we fit linear mixed-effect models, using each plateau as a random effect to account for site-specific characteristics and spatial autocorrelation. In the case of *B. carajensis* from Serra Norte, we also included a random effect specifying the genetic cluster containing each individual (see genetic structure results using the Admixture software). In Serra Sul, which comprises a single large plateau, we used generalized least-squares models (GLS) fitted with different correlation structures (linear, exponential, Gaussian, and spherical) to explicitly model spatial autocorrelation. The “*weight*” argument was used in some cases to account for heteroscedasticity. Raw *f* and logit-transformed *H* were used as response variables and models fitted using the *nmle* R package (Pinheiro et al., 2009). For each model, we calculated AIC, the difference of each model and the best model (ΔAIC) and the Akaike’s weight of evidence (wAIC). Models with ΔAIC ≤ 2 were considered as equally plausible (Zuur et al., 2009). The set of best models (ΔAIC ≤ 2) were compared to reduced models without each predictor variable, using likelihood ratio tests (LRT, *α* = 0.05), and all models were validated by plotting residual *vs.* fitted values and by checking for residual autocorrelation. Relative variable importance was calculated summing the Akaike weights of the best-fitting models in which the variable of interest was present (https://github.com/carolinacarvalho/Importance_plot). Model averaging across the set of best models was used to compute parameters estimates that account for uncertainty in model selection (Burnham and Anderson, 2002). We also estimated confidence intervals for parameter estimates to assess the statistical power of our models (Hoenig and Heisey, 2001).

### Gene flow

To assess the effect of habitat loss on gene flow, we first optimized gene flow hypotheses and then tested them by modeling isolation by resistance (IBR, (McRae, 2006)). Yang’s genetic relatedness between pairs of individuals (Yang et al., 2010) was used as a proxy for recent gene flow, given that it represents the number of common ancestors in the recent past (Wang, 2017). Although relatedness is not a direct measure of gene flow, it has been widely used to describe genetic connectivity among individuals (Balkenhol et al., 2016; Monteiro et al., 2019; Storfer et al., 2010) and a recent simulation study showed that it is among the most accurate individual-based genetic distance metric for landscape genetic studies (Shirk et al., 2017). Resistance to gene flow due to mining was modeled using land cover maps for different years (2016, 2014, 2011 and 1979) containing only the major land cover classes of our study region: Montane Savanna (Canga), Forest (evergreen forest) and Mine. Water bodies, Pasture and Urban areas were excluded because they occurred outside the extent of our samples (Fig. 1). By so doing we were able to evaluate the permeability to gene flow of each land cover class; and test whether habitat loss driven by mining hindered gene flow across our replicated landscapes. Additional variables found to be important predictors of gene flow in other plants (Dyer, 2016; Lanes et al., 2018) were modeled along with land cover, including geographic distance, elevation (DEM retrieved from the USGS Earth Explorer), terrain roughness (generated from the DEM using the Terrain Analysis plug-in from QGIS), and bioclimatic variables (retrieved from WorldClim). To select a set of orthogonal variables explaining most climatic variation across our study area, we first ran separate principal component analyses (PCA) for each species using the extracted values from all 19 WorldClim bioclimatic layers plus elevation (scaled) (Fig. S2). We then selected the three variables showing the strongest correlation with the first, second and third PCA axis (which explained more than 85% of total variance in both *B. carajensis* and *M. carajensis*). These were minimum temperature of coldest month (bio06) and precipitation of wettest (bio16) and coldest quarter (bio19) for *B. carajensis*; and minimum temperature of coldest month (bio06), precipitation of wettest quarter (bio16) and temperature seasonality (bio04) for *M. carajensis*.

A genetic algorithm (unrelated to the genetic data) was implemented through the *ResistanceGA* package to generate optimized resistance surfaces for each one of these variables (Peterman, 2018). The advantage of this optimization procedure is that it relies on empirical genetic data, ensuring that resistance values attributed to resistance surfaces will relate meaningfully to the movement of genes across the landscape (Peterman, 2018). Moreover, since resistance values are not defined *a priori*, the optimized resistance surfaces can be considered unbiased by any existing knowledge or human preferences. In the case of land cover maps, random initial resistance values were assigned for each class; then pairwise resistance distances were measured using random-walk commute times; and finally pairwise genetic distance was regressed on resistance distance using maximum likelihood population effect models (MLPE, see below). The whole process was iterated until no significant change was found in the objective function (Peterman, 2018). We then performed the same steps for the remaining continuous predictors, but instead of assigning random initial resistance values, eight types of transformations were applied to the raw values. In this case, two parameters controlling Ricker and Monomolecular functions were iteratively varied during the optimization (Peterman, 2018). Ten independent runs of optimization were conducted for each surface to assess the convergence in parameter estimates (Khimoun et al., 2017). All rasters were set to Universal Transverse Mercator (UTM) projection, and cropped to the extent of sampling locations plus a buffer area of 5 km to minimize border effects (Lanes et al., 2018). Land cover resistance surfaces and terrain roughness were optimized using 250 × 250 m resolution maps, while 900 × 900 m resolution maps were used for WorldClim layers as this is the highest available. Serra Norte and Serra Sul were analyzed separately aiming to replicate IBR analyses in two separate areas exposed to open-pit mining.

Using the program Circuitscape V4.0 (McRae, 2006), we then calculated pairwise resistance distances between all samples, employing the optimized resistance surfaces described above plus a surface where all pixels were set to 1 to create a null model of isolation by geographic distance (IBD). To assess isolation by resistance (IBR), defined as the correlation between genetic and resistance distances (McRae, 2006), we fitted mixed-effects regression models using penalized least squares and a correlation structure designed to account for the non-independence of pairwise distances (maximum-likelihood population effects - MLPE: https://github.com/nspope/corMLPE; (Clarke et al., 2002)). Yang’s genetic relatedness between individuals was used as the response variable and the different resistance distances (contemporary and historical land cover, elevation, terrain roughness, temperature, precipitation, and geographic distance) as predictors. All MLPE models accounted for the underlying population structure, either by considering only individuals belonging to the same genetic cluster (most cases), or by including an additional random effect specifying if pairwise distances represented individuals from the same or from different genetic clusters (the case of *B. carajensis* from Serra Norte, see genetic structure results using the Admixture software). To evaluate the incidence of time-lag effects potentially masking mining effects on gene flow, we first fitted uni-variate models for each species and region using resistance distances from land cover surfaces from all years, plus those from geographic distance surfaces. The best models were selected using the Akaike Information Criterion (ΔAIC < 2), and whenever geographic distance was found among the best models we considered IBD as the most parsimonious gene flow model (Balkenhol et al., 2016; Burnham and Anderson, 2002). To evaluate the sensitivity of our analysis to the resolution (grain size) of spatial data, we also compared uni-variate land cover models containing resistance distances computed from surfaces with different grain sizes (100 × 100 m, 300 × 300 m, 600 × 600 m and 900 × 900 m). Results were consistent across the different resolutions (Table S2), so we ran all subsequent analysis using a grain size of 900 × 900 m. We then fitted multiple regression models containing resistance distances from the best uni-variate land cover models selected in the previous step and resistance distance from all other optimized surfaces for each species and region. Models containing all possible combinations of non-collinear predictors (*r* < 0.6, Fig. S3) were compared using the *dredge* function from the package *MuMIn* (https://github.com/rojaff/dredge_mc; (Barton and Barton, 2015)), and best models were selected using AIC. Likelihood ratio tests (LRT) were performed to assess the influence of each predictor variable on the best model’s log-likelihood (Jaffé et al., 2016), and relative variable importance, model-averaged parameter estimates and confidence intervals were calculated as described above. Finally, we carried out a barrier analysis to identify genetic discontinuities between individuals by using Monmonier’s algorithm and Gabriel’s graph implemented in package *adegenet* (Jombart and Ahmed, 2011).

### Germination experiments

To evaluate if seeds from both study species are able to germinate inside iron ore mines, we ran a set of germination experiments. Seeds from both species were sown over four different substrates (Whatman® paper, Canga topsoil, forest topsoil, and mining waste substrate) placed in plastic boxes (Gerbox – 11 × 11 × 4 cm) and kept in a growth chamber (Fitotron SGC 120, Weiss Technik, UK) under continuous darkness, constant temperature (20°C) and air humidity (60%) for 33 consecutive days, from September 4th to October 7th 2018. Substrates received distilled water until the retention capacity, and water losses by evaporation were replaced daily. All treatments were carried out with five replicates for each substrate in each species. Each replicate contained 25 seeds from *B. carajensis* and 12 seeds from *M. carajensis*. The number of germinated seeds was recorded daily, with germination defined as the emission of 2 mm of primary root.

## Results

### Neutral dataset

We collected leaf tissue samples of 150 individuals of *B. carajensis* and 207 individuals of *M. carajensis* distributed across the entire occurrence range of both species and surrounding two large iron ore mines (Fig. 1). Samples were frozen and their DNA later extracted and shipped for genotyping-by-sequencing (RAD-sequencing) and bioinformatic processing. We identified a total of 10,016 SNPs in *B. carajensis* and 20,464 SNPs in *M. carajensis*, but after filtering these for missing data, quality, depth, linkage disequilibrium, deviations from the Hardy-Wenberg Equilibrium and *F*_*ST*_ outlier loci, we obtained sets of neutral and independent markers containing 1,411 and 6,052 loci for each species, respectively.

### Genetic structure

Two complementary genetic clustering approaches used to assess population structure (Admixture and DAPC) indicated the presence of three clusters in *B. carajensis* and two in *M. carajensis* (Fig. 2, Fig. S4-S6). Cluster-level heterozygosity was slightly higher in *B. carajensis* than in *M. carajensis*, and significant albeit low inbreeding was found in one genetic cluster of each species (Fig. 2). Both species showed spatial autocorrelation in genetic relatedness in Serra Norte but not in Serra Sul, and the strength of spatial autocorrelation was higher in *B. carajensis* (Fig. 2).

**Fig. 2.**
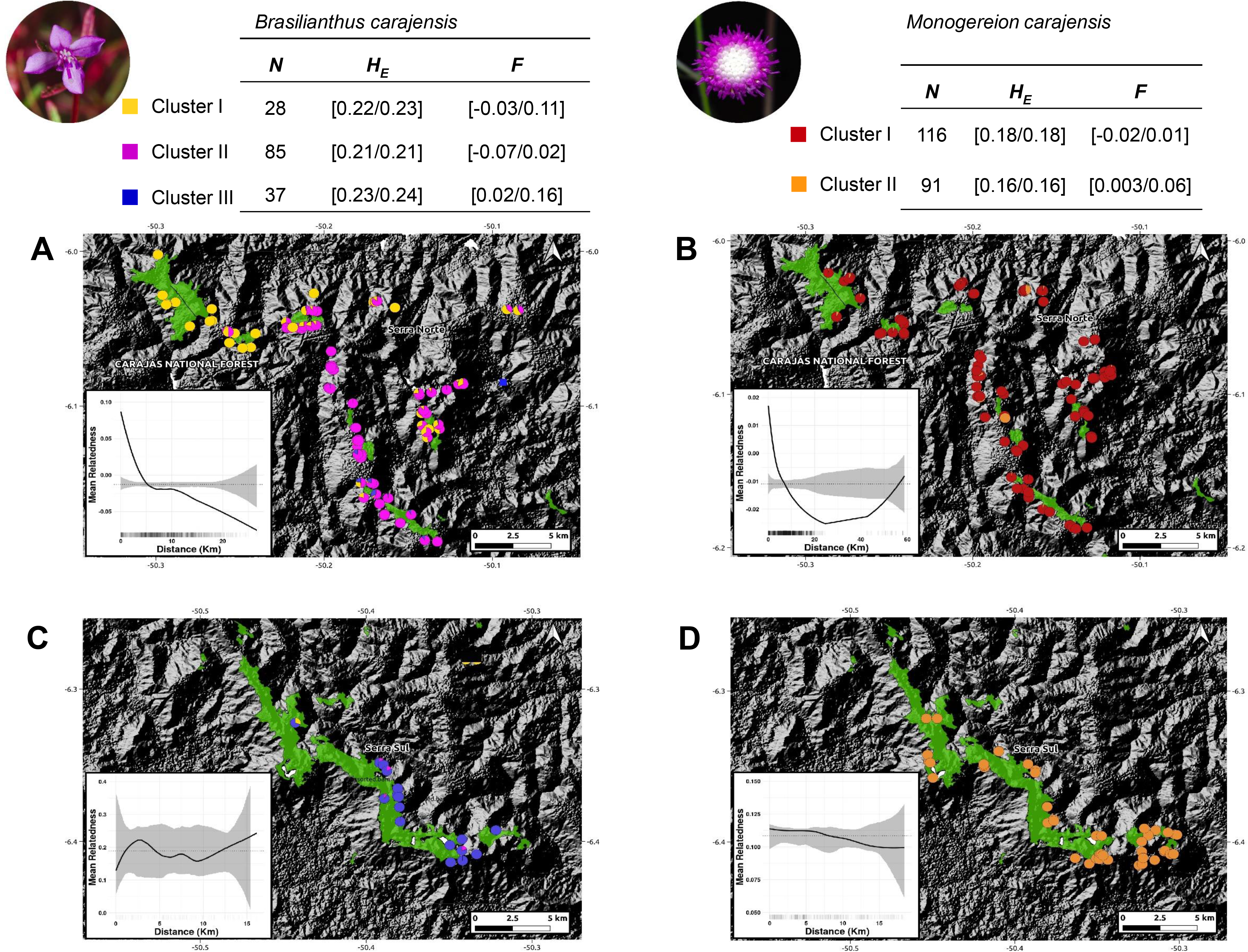
Map showing the ancestry coefficients from *Brasilianthus carajensis* (A and C) and *Monogereion carajensis* (B and D) in Serra Norte (upper panels) and Serra Sul (lower panels) determined using the Admixture software. Montane Savanna areas are shown in green against hill shade layers. Smaller lower-left corner plots show spatial autocorrelation in genetic relatedness, where black solid lines are the LOESS fit to the observed relatedness, gray shaded regions are 95% confidence bounds around the null expectation (black dotted lines), and short vertical lines at the bottom of the figure are observed pairwise distances. Genetic diversity measures for each genetic cluster are shown in the upper tables. Sample sizes (N) are followed by mean expected heterozygosity (*H*_*E*_) and mean inbreeding coefficient (*F*), and values represent 95% confidence intervals.

### Genetic diversity

To assess the effect of habitat loss on genetic diversity, we regressed individual-level diversity metrics on historical habitat amount and habitat loss driven by mining in different years. Heterozygosity (*H*) and inbreeding (*f*) were not influenced by habitat loss, either in Serra Norte nor in Serra Sul, as the set of best-fitting models always included null models or historical (pre-mining) habitat amount (Fig. 3, Table S3). Although confidence intervals for the effect of habitat loss on heterozygosity were usually narrow, those describing the effect of habitat loss on inbreeding were very broad in *B. carajensis* and moderately so in *M. carajensis* (Table S4). Historical habitat amount was found to be associated with inbreeding in both species, although the direction of the effect varied (Fig. 4, Table S4).

**Fig. 3.**
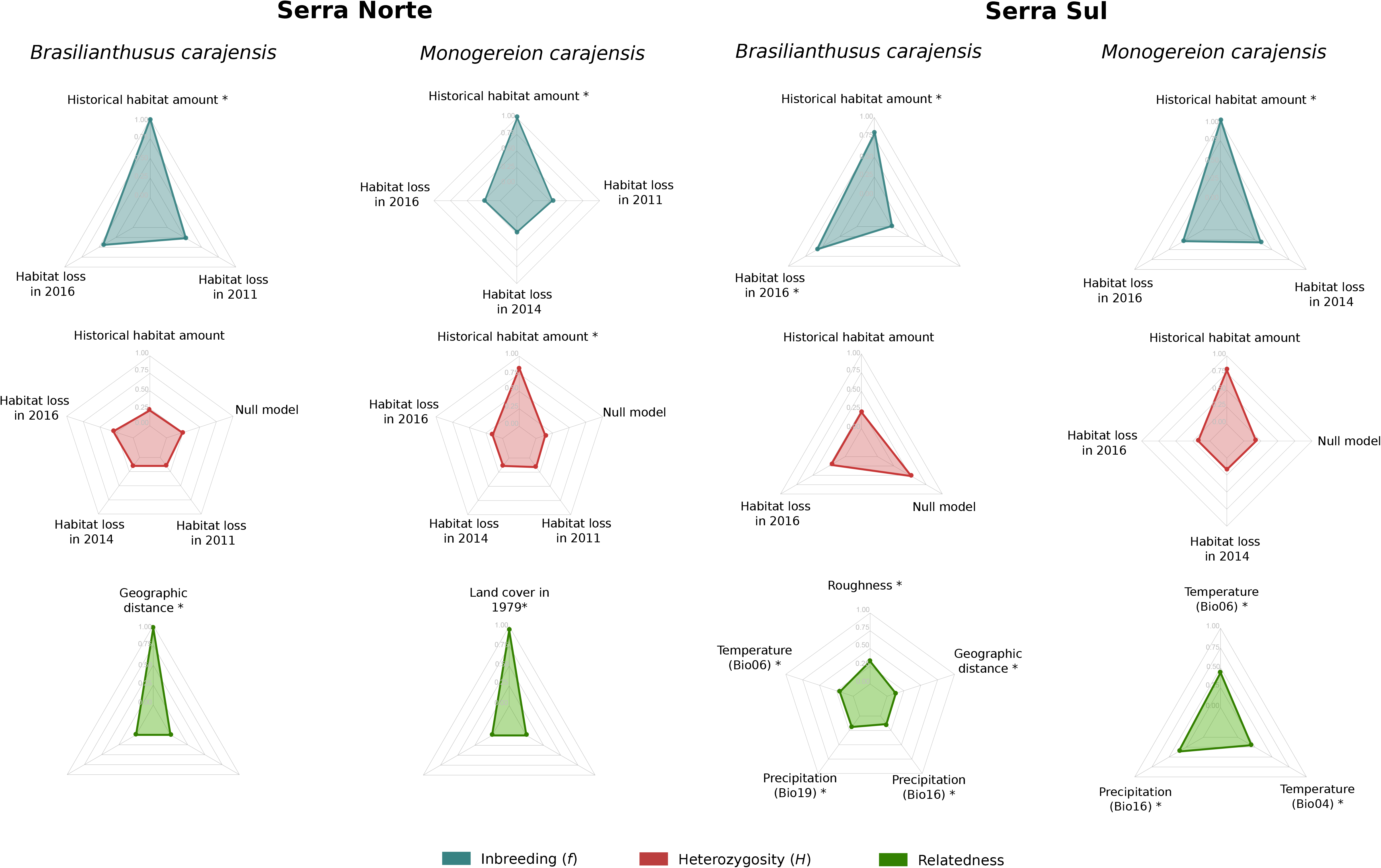
Relative variable importance in the set of best-fitting models (ΔAIC ≤ 2) for *Brasilianthus carajensis* and *Monogereion carajensis* in Serra Norte and Serra Sul (see methods and Supplementary Tables S3 and S5 for details). Individual-level genetic diversity metrics (*H* and *f*) were response variables and habitat amount in 1979 and habitat loss in 2011, 2014 and 2016 were predictors in genetic diversity models. Pairwise inter-individual genetic relatedness was the response variable and resistance distances computed from optimized surfaces were predictors in IBR models. Likelihood Ratio Test (LRT) were performed to assess if each predictor variable significantly improved the model’s log-likelihood (significance levels are highlighted with: *p < 0.05; **p < 0.01; and ***p < 0.001).

**Fig. 4.**
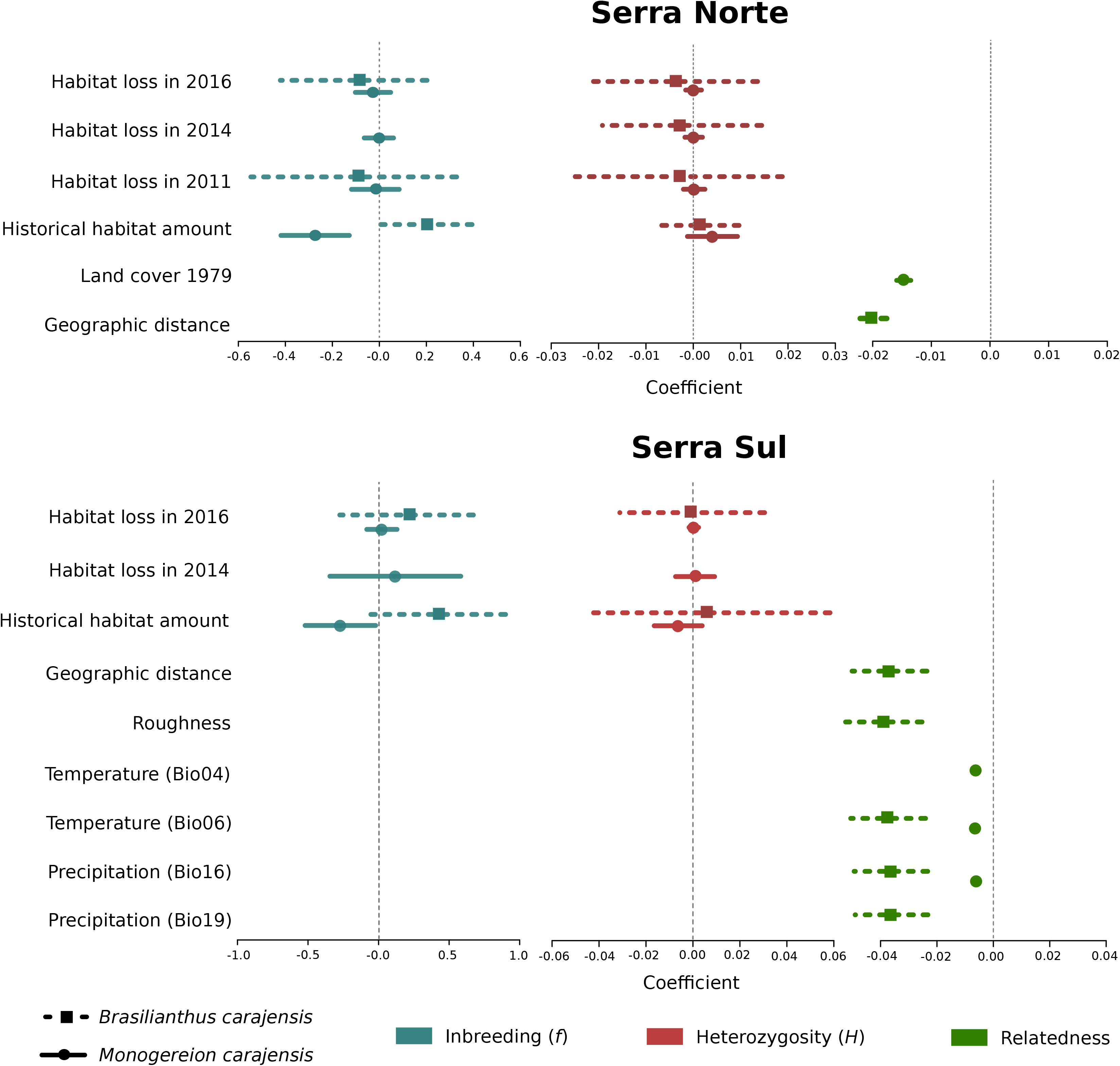
Coefficient plots for the set of best-fitting models (ΔAIC ≤ 2) for *Brasilianthus carajensis* and *Monogereion carajensis* in Serra Norte and Serra Sul (see methods and Supplementary Tables S4 and S6 for details). Points represent model-averaged regression coefficients and lines the 95% confidence intervals.

### Gene flow

To assess the effect of habitat loss on gene flow, we first employed a genetic algorithm to optimize gene flow hypotheses, then calculated resistance distances between individual samples, and finally modeled isolation by resistance (IBR) regressing pairwise genetic relatedness on resistance distances through Maximum Likelihood Population Effects (MLPE) models. While resistance to gene flow due to mining was modeled using land cover maps for different years (2016, 2014, 2011 and 1979), additional covariates modeled along with land cover included geographic distance, terrain roughness, elevation, and bioclimatic variables. The optimization of resistance surfaces revealed that Canga was the land cover class representing lowest resistance to gene flow in both species, whereas mining areas and evergreen forests imposed higher resistance (Fig. S7-S10). However, univariate MLPE regression models revealed that geographic distance usually explained relatedness patterns as well as land cover (Table S2), and only pre-mining land cover (1979) was found to explain relatedness patterns better than geographic distance in *M. carajensis* from Serra Norte. Our results thus reveal that mining neither hinders nor facilitates gene flow in these two endemic annual plants. While these results hold across different resolutions (Table S2), an independent barrier analysis also failed to identify barriers between individuals separated by mining areas (Fig. S11). Multiple MLPE regression models showed that isolation by geographic distance (IBD) explained genetic relatedness patterns in *B. carajensis*, whereas IBR was more important in *M. carajensis* (Fig. 3, Table S5). In all cases, genetic relatedness decreased with increasing resistance (Fig. 4, Table S6).

### Germination experiments

Germination experiments revealed that seeds from both species are able to germinate in mining waste substrates. Whereas *M. carajensis* showed similar germination in Canga and mining substrates, germination rates of *B. carajensis* were higher in Canga topsoil (Fig S12).

## Discussion

Our study is the first to assess the genetic consequences of habitat loss while accounting for all the major limitations constraining the quantification of habitat amount effects on genetic variation. Our results reveal that habitat loss driven by mining did not affect genetic diversity or gene flow in two endemic herbs from Amazonian Savannas. Whereas historical habitat amount was found to influence inbreeding, heterozygosity and inbreeding were not affected by habitat loss in either species. Finally, gene flow was mainly influenced by geographic distance in *B. carajensis* and by pre-mining land cover and local climate in *M. carajensis*.

The genetic structure in *B. carajensis* mirrored that from the co-occurring perennial morning glory *Ipomoea maurandioides* (Lanes et al., 2018), showing two differentiated genetic clusters in Serra Norte, while *M. carajensis* only presented one cluster in Serra Norte and another one across the remaining distribution range. This genetic structure was considered when assessing landscape effects on genetic diversity and gene flow in order to control for historical demographic processes. Additionally, since open-pit mining in our study region did not usually result in decreased structural connectivity between habitat patches (since these were already separated, Fig. S1), we were able to assess habitat loss effects that were not heavily influenced by habitat fragmentation (Fahrig, 2003).

The maintenance of genetic diversity in spite of extreme habitat loss suggests that our study plants are able to colonize mining environments and maintain gene flow across open-pit mines. Germination experiments revealed that seeds from both species can indeed germinate in mining waste substrates. Additionally, both plant species showed extensive gene flow across mining areas, and mining neither enhanced nor hindered gene flow. Similar results were found for a threatened orchid and the American pika, which showed analogous levels of genetic diversity in mining and natural habitats (Esfeld et al., 2008; Waterhouse et al., 2017), although neither gene flow nor historical effects were assessed. Inbreeding levels in our focus species are comparable to those observed in the widespread *I. maurandioides* (Lanes et al., 2018), and since they were associated with historical habitat amount they seem to reflect density-dependent selfing (Leimu et al., 2006). Our results thus reveal that some insect-pollinated and wind-dispersed plants do not experience genetic erosion due to habitat loss (Aguilar et al., 2008, 2019). A possible mechanism explaining the maintenance of genetic diversity is seed dormancy over long periods of time, which would result in multiple overlapping generations being represented in the seed bank (Honnay et al., 2008).

Both species presented spatial autocorrelation in genetic relatedness in Serra Norte but not in Serra Sul, indicating a more restricted gene flow in the Canga archipelago of Serra Norte than in the large continuous plateau of Serra Sul. Additionally, geographic distance was weakly correlated with recent land cover resistance in Serra Norte but not in Serra Sul, where it was strongly correlated with land cover resistance from all years (Fig. S3). We thus expected that isolation by resistance (IBR) would be easier to disentangle from isolation by distance (IBD) in Serra Norte than in Serra Sul. In Serra Norte, however, geographic distance and pre-mining land cover (highly correlated with geographic distance) were the best predictors of current gene flow in *B. carajensis* and *M. carajensis* populations, respectively. Considering the strong winds characterizing Montane Savanna ecosystems from the Carajás Mineral Province (Skirycz et al., 2014), and the fact that wind currents in open landscapes are known to facilitate long-distance dispersal of plant propagules (Heydel et al., 2014; Soons et al., 2004), we posit that wind-mediated dispersal is driving gene flow across Montane Savannas and open-pit mines. High levels of gene flow have also been detected in others wind-dispersed species in open anthropogenic landscapes like agricultural areas (Aavik et al., 2013; Heydel et al., 2014; Kamm et al., 2010), suggesting that open areas promote genetic connectivity in wind-dispersed plants. On the other hand, local climate differences also appear to explain gene flow patterns in *M. carajensis* populations from Serra Sul better than IBD, suggesting mismatches in flowering periods (Dick et al., 2008) or different local adaptations (Hoffmann and Sgrò, 2011; Lenormand, 2002). We nevertheless caution that our study design and the little available knowledge on the natural history of these plants do not allow disentangling the relative contribution of pollen and seed dispersal on gene flow.

The absence of an effect of habitat loss on genetic variation can be attributed to time-lags between the onset of disturbances and genetic responses (Schlaepfer et al., 2018). We overcame this methodological limitation by focusing on species with a short generation time (i.e. completing their life cycle within one year), and by explicitly incorporating time scale into our analyses (evaluating land cover maps from different years). Moreover, historical demographic processes are unlikely to have biased our results, since our isolation by resistance models explicitly accounted for the underlying population structure. Mining operations began in the 1980s in Serra Norte, allowing enough time (~40 generations) to assess genetic responses to mining. On the other hand, Serra Sul was still pristine by 2013, so only three plant generations were exposed to mining before our samples were collected in 2017. This could explain why in Serra Sul recent land cover did not explain relatedness patterns better than geographic distance alone (Table S2). However, the fact that recent land cover did not explain relatedness patterns in either species in Serra Norte, strongly suggests that gene flow has been maintained across mines. In contrast, land cover in existence two decades ago was found to explain gene flow in a perennial narrow endemic morning glory occurring in Serra Norte (Lanes et al., 2018), indicating that our methods should be sufficient to detect an effect of mining should there be one, although differences in reproductive systems and dispersal modes could also underlie these different results (Aguilar et al., 2008; Vranckx et al., 2012). Additionally, our findings were unaffected by the resolution of spatial data and were supported by an independent barrier analysis, so they strongly indicate that gene flow in our two annual herbs is unaffected by habitat loss driven by mining.

The incidence of time-lag effects on the response of genetic diversity to habitat loss is nevertheless more difficult to assess, since different metrics respond at different rates. Empirical and simulations studies have shown changes in inbreeding and allelic richness immediately after the onset of disturbances, whereas heterozygosity is usually lost more slowly, over subsequent generations (Keyghobadi et al., 2005; Lowe et al., 2005). We therefore caution that longer time lags would be needed to rule out an effect of habitat loss on the observed heterozigosity of our study species. On the other hand, ~40 generations should be enough to detect a response in the levels of inbreeding, and confidence intervals for the effect of habitat loss on inbreeding (Table S4) suggest that our models had sufficient power to identify non-significant effects. Our results thus indicate, with moderate confidence, that habitat loss did not result in increased inbreeding in our study plants.

## Conclusions

Using thousands of genetic markers to study two annual endemic plants in replicated landscapes, we found that extreme habitat loss driven by mining did not result in any detectable genetic consequences. Since our results are largely unbiased by the effect of habitat fragmentation, the underlying genetic structure of plant populations, the resolution of spatial data, or time-lag effects, they reveal that habitat loss does not always entail negative genetic consequences. Although habitat fragmentation has been shown to disrupt gene flow and increase inbreeding across plants species, regardless of their characteristics (Aguilar et al., 2019), our study unveils remarkably resilient species to extreme habitat loss as similar levels of genetic diversity and gene flow were found in mining and natural habitats. These findings imply that it is not possible to generalize about the genetic consequences of habitat loss, so future conservation efforts need to consider species individually.

## Acknowledgments

Funding was provided by Instituto Tecnológico Vale, Conselho Nacional de Desenvolvimento Científico e Tecnológico (CNPq) grants 301616/2017-5 (RJ), 307479/2016-1 and 402756/2018-5 (GO), and 300714/2017-3 (EL), and Coordenação de Aperfeiçoamento de Pessoal de Nível Superior (CAPES) grant 88887.156652/2017-00 (CSC). We thank Alexandre Castilho, Cesar Neto and Waléria Monteiro for assistance in the field, Gleiciane Salvador and Manoel Lopes for help in the laboratory, Prof. Ing. Jaroslay Dolezel for provinding the standards used in flow cytometry analyses, and Nathaniel Pope for advice on the statistical analyses. This manuscript has been released as a Pre-Print at bioRxiv (Carvalho et al., 2019).

## Author contributions

RJ conceived, designed and coordinated the project. RJ and PLV coordinated the field work and sampling. ECML, ARS, CFC and MG performed laboratory work. RJ, ECML, CSC, ARS and CFC performed the data analysis. The first draft of the paper was written by CSC and ECML with input from RJ. All authors contributed to discussing the results and editing the paper.

## Data Availability Statement

Geographic coordinates in decimal degrees, genotypes in Variant Call Format and sequences in FASTA format for both species are provided here: https://figshare.com/articles/Habitat_loss_does_not_always_entail_negative_genetic_consequences/8224175. All the mentioned R scripts have been deposited in GitHub and their url addresses provided in the text.

