## Supporting Information for "Habitat loss does not always entail negative genetic consequences"

### *Supplementary Material*

#### Supplementary Figures

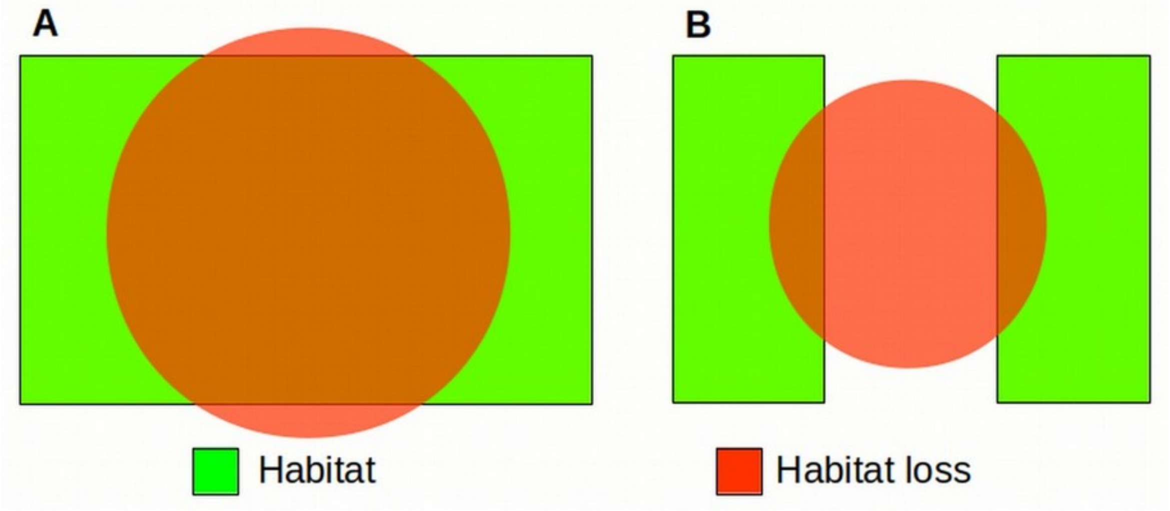


**Figure S1:** Habitat loss with fragmentation (A) and without fragmentation (B). In the later case habitat loss does not result in decreased structural connectivity between habitat patches, since these were already separated. This is the dominant pattern of habitat loss resulting from open-pit mining in our study region (Fig. 1).

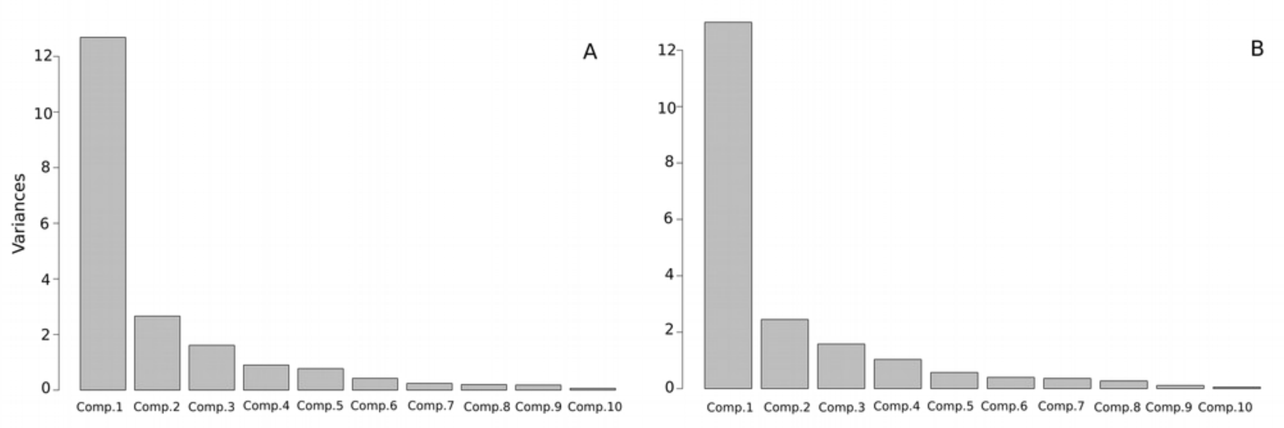


**Figure S2:** Scree plot of the first ten principal components summarizing bioclimatic variables and elevation for *Brasilianthus carajensis* (A) and *Monogereion carajensis* (B).

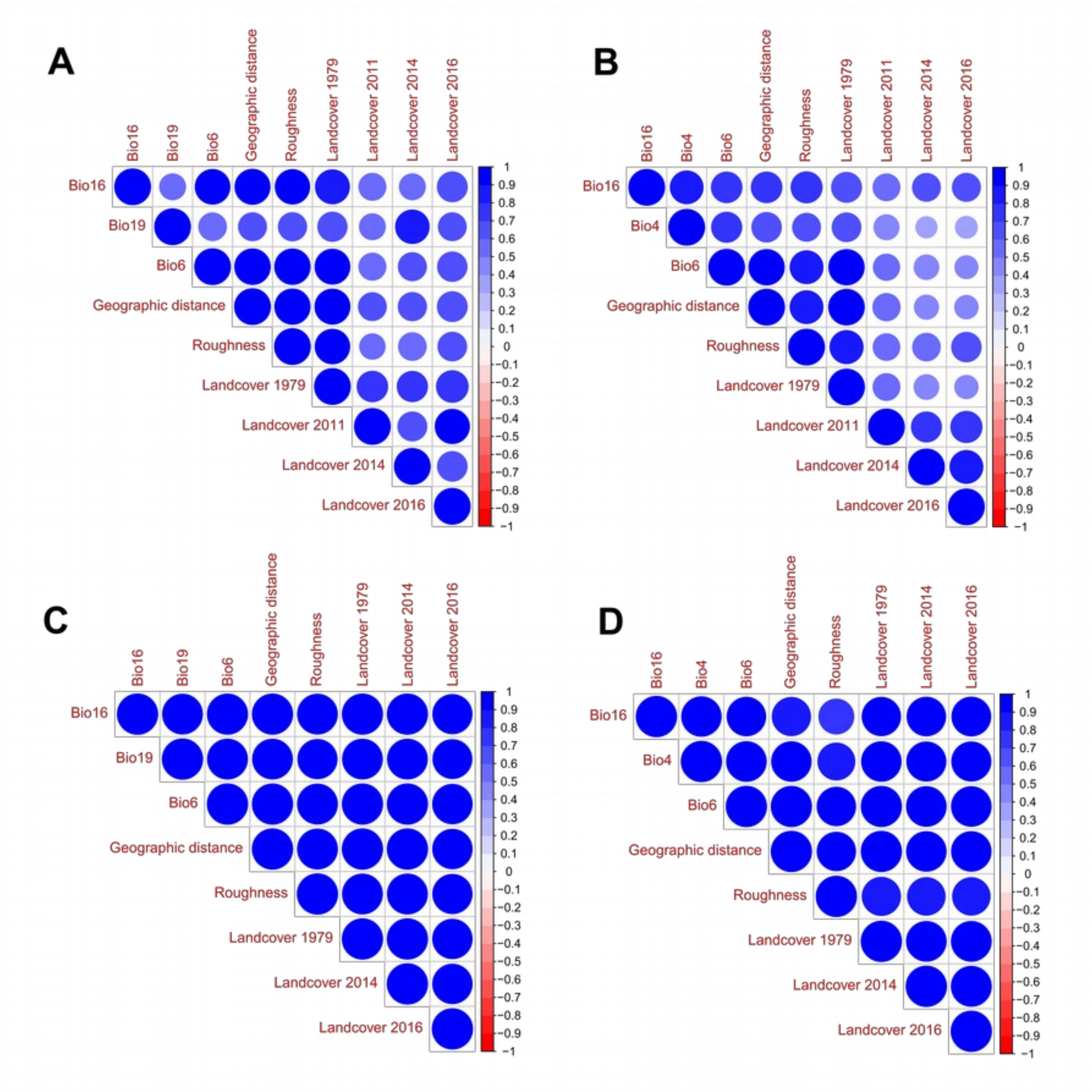


**Figure S3:** Pearson's correlation among resistance distance pairs (land cover for year 1979, 2011, 2014 and 2016; terrain roughness, temperature - Bio6 and Bio4, precipitation - Bio19 and Bio16; and geographic distance) for *Brasilianthus carajensis* (A and C) and *Monogereion carajensis* (B and D) individuals in Serra Norte (upper panels) and Serra Sul (lower panels).

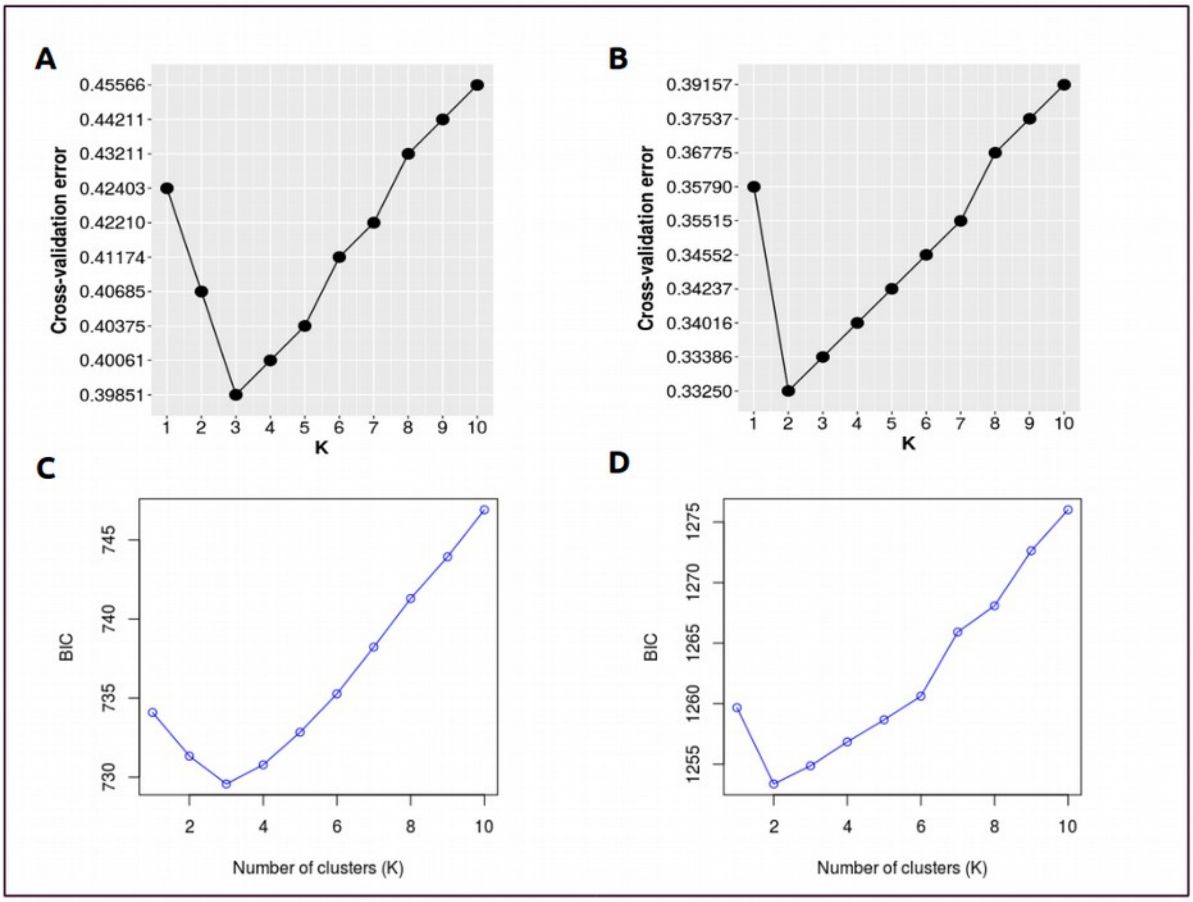


**Figure S4:** Plots showing the optimal number of genetic clusters (k) for *Brasilianthus carajensis* (A and C) and *Monogereion carajensis* (B and D) based on the Admixture program (upper panels) and on the discriminant analysis of principal components – DAPC (lower panels). In the Admixture program the optimal k is based on cross-validation errors. In the DAPC the minimum value of Bayesian Information Criterion (BIC) indicate best-supported number of genetic cluster.

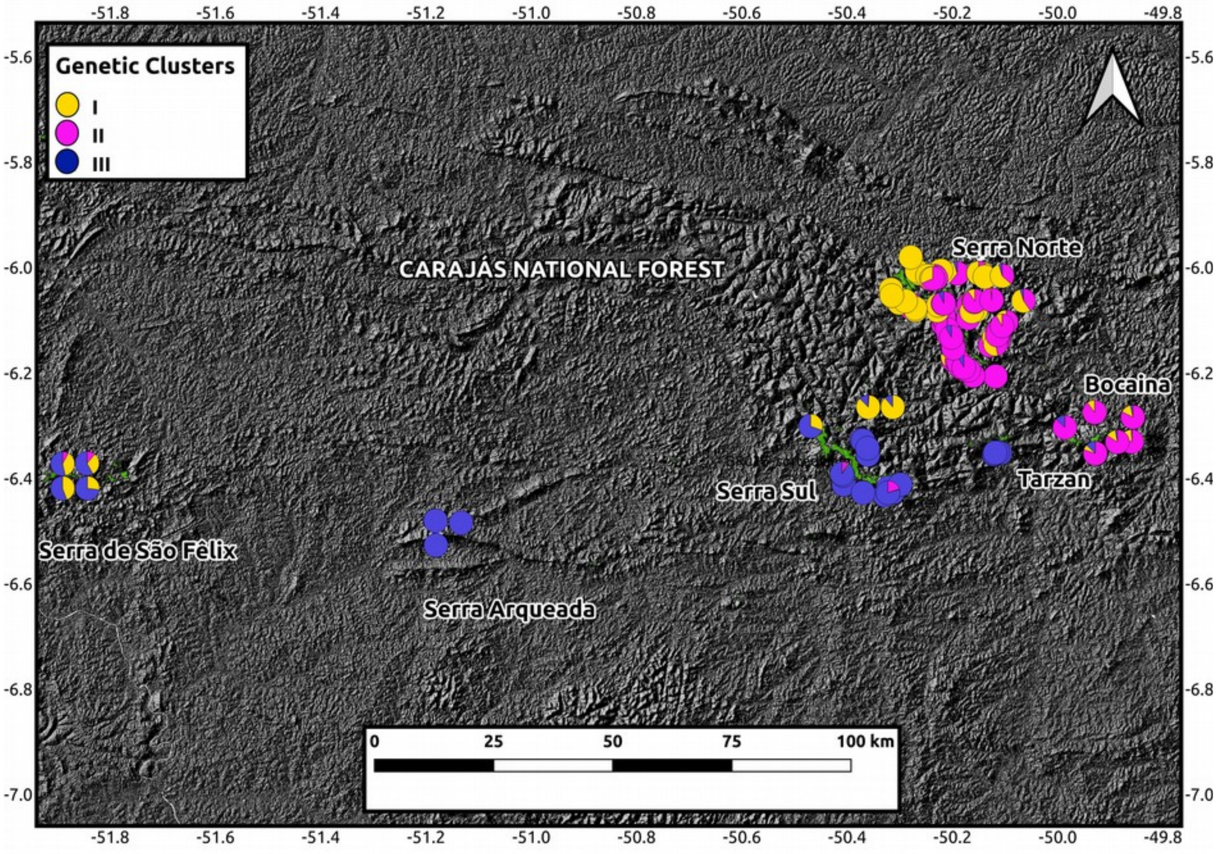


**Figure S5:** Map showing *Brasilianthus carajensis* ancestry coefficients determined using Admixture. Montane Savanna areas are shown in green against an elevation (hill shade) layer.

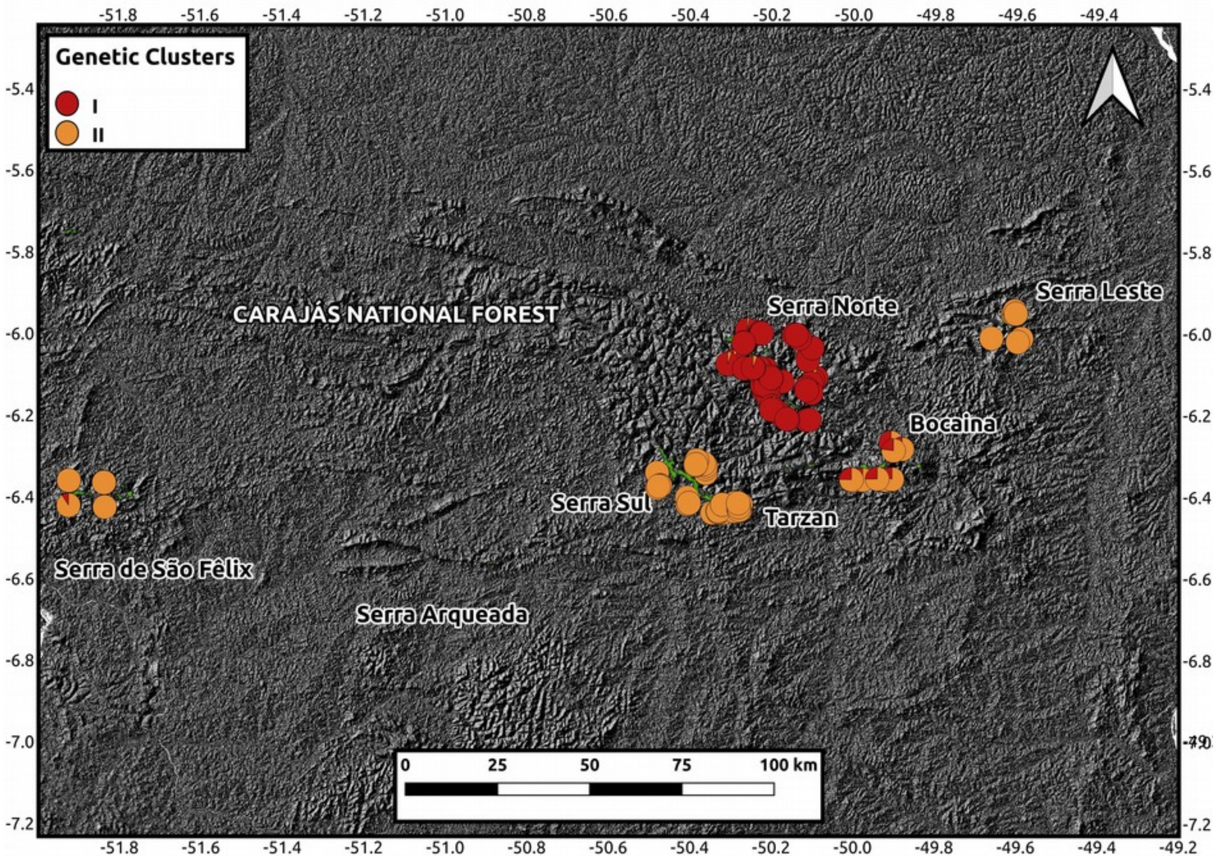


**Figure S6:** Map showing *Monogereion carajensis* ancestry coefficients determined using Admixture. Montane Savanna areas are shown in green against an elevation (hill shade) layer.

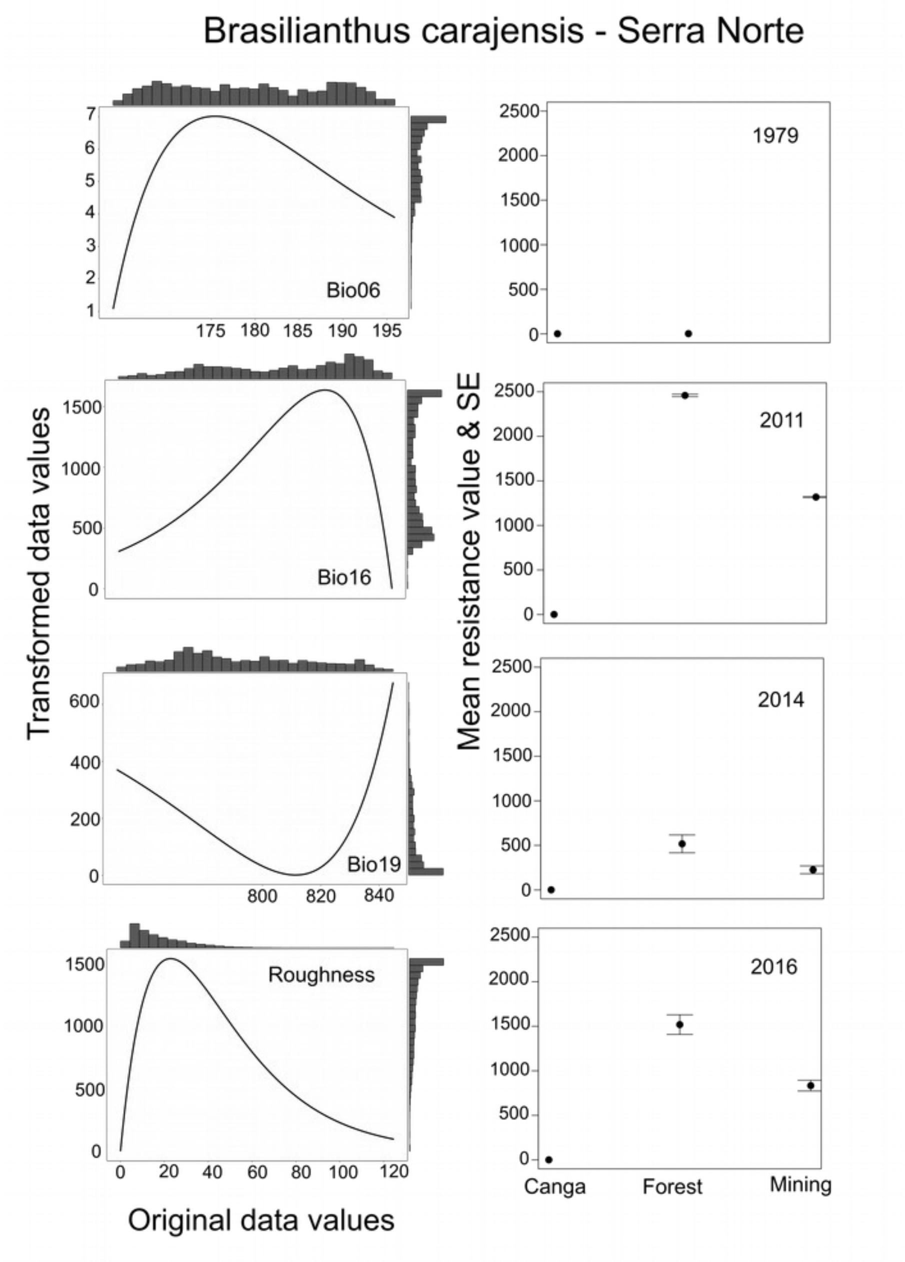


**Figure S7:** Transformation applied to each of continuous (left panels) and categorical (right panels) variables to generate resistance surfaces for *Brasilianthus carajensis* from Serra Norte. For continuous variables, original data values are represented in x-axes, while transformed data values are shown in y-axes, and histograms present the distribution of original untransformed and transformed variables. For categorical variables, land cover classes were averaged across ten replicates and standard errors (SE) are provided. Categorical resistance surfaces were built for different years (1979, 2011, 2014 and 2016). Bio06 represents the minimum temperature of the coldest month, and Bio16 and Bio19 represent precipitation of wettest and coldest quarter, respectively.

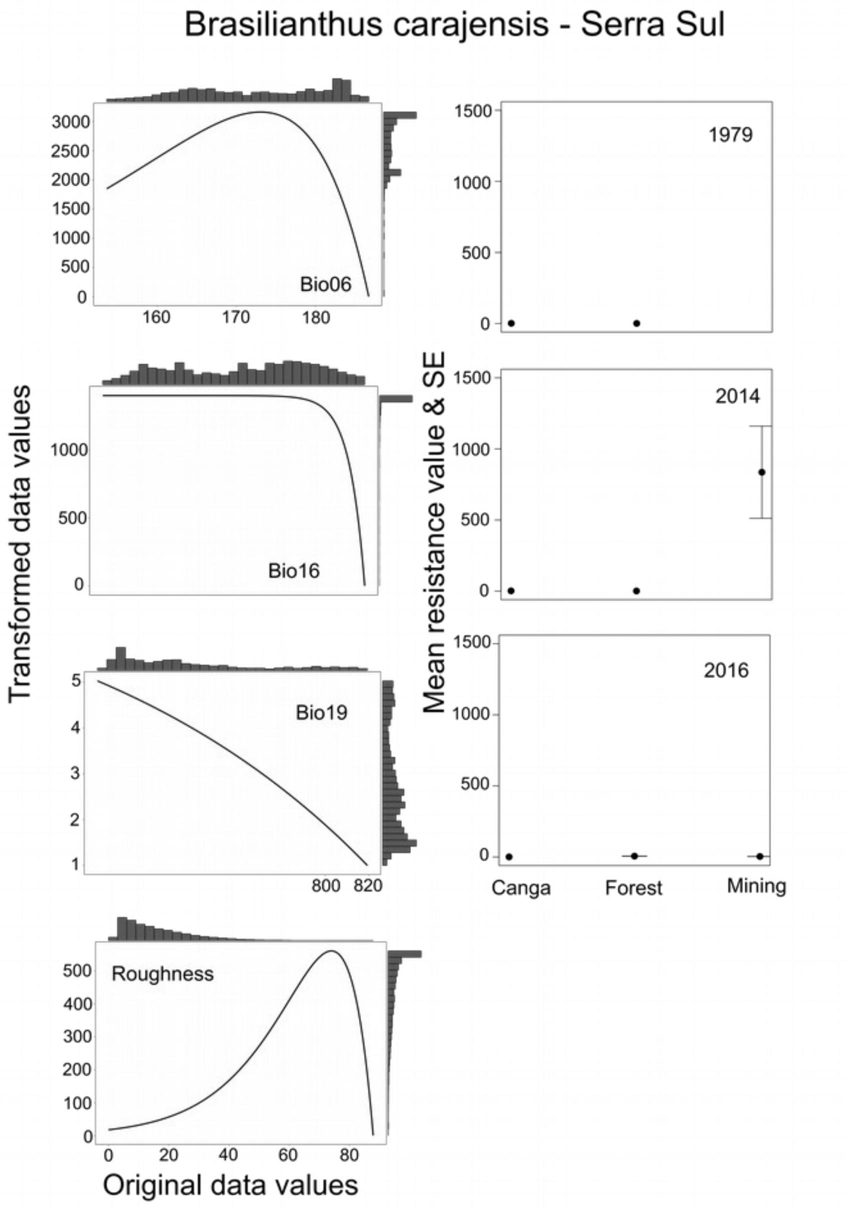


**Figure S8:** Transformation applied to each of continuous (left panels) and categorical (right panels) variables to generate resistance surfaces for *Brasilianthus carajensis* from Serra Sul. For continuous variables, original data values are represented in x-axes, while transformed data values are shown in y-axes, and histograms present the distribution of original untransformed and transformed variables. For categorical variables, land cover classes were averaged across ten replicates and standard errors (SE) are provided. Categorical resistance surfaces were built for different years (1979, 2014 and 2016). Bio06 represents the minimum temperature of the coldest month, and Bio16 and Bio19 represent precipitation of wettest and coldest quarter, respectively.

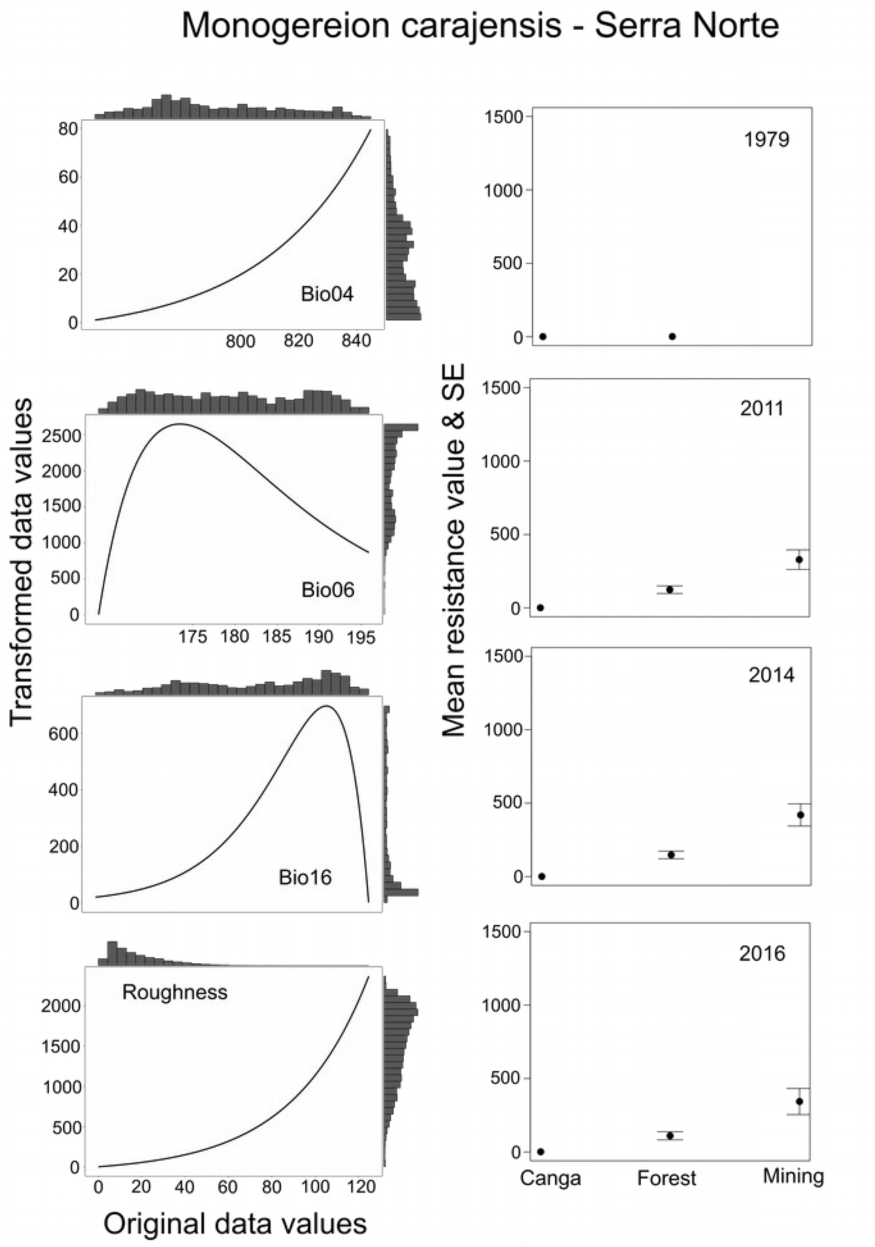


**Figure S9:** Transformation applied to each of continuous (left panels) and categorical (right panels) variables to generate resistance surfaces for *Monogereion carajensis* from Serra Norte. For continuous variables, original data values are represented in x-axes, while transformed data values are shown in y-axes, and histograms present the distribution of original untransformed and transformed variables. For categorical variables, land cover classes were averaged across ten replicates and standard errors (SE) are provided. Categorical resistance surfaces were built for different years (1979, 2011, 2014 and 2016). Bio04 and Bio06 represent temperature seasonality and the minimum temperature of the coldest month, respectively, while Bio16 signifies precipitation of wettest quarter.

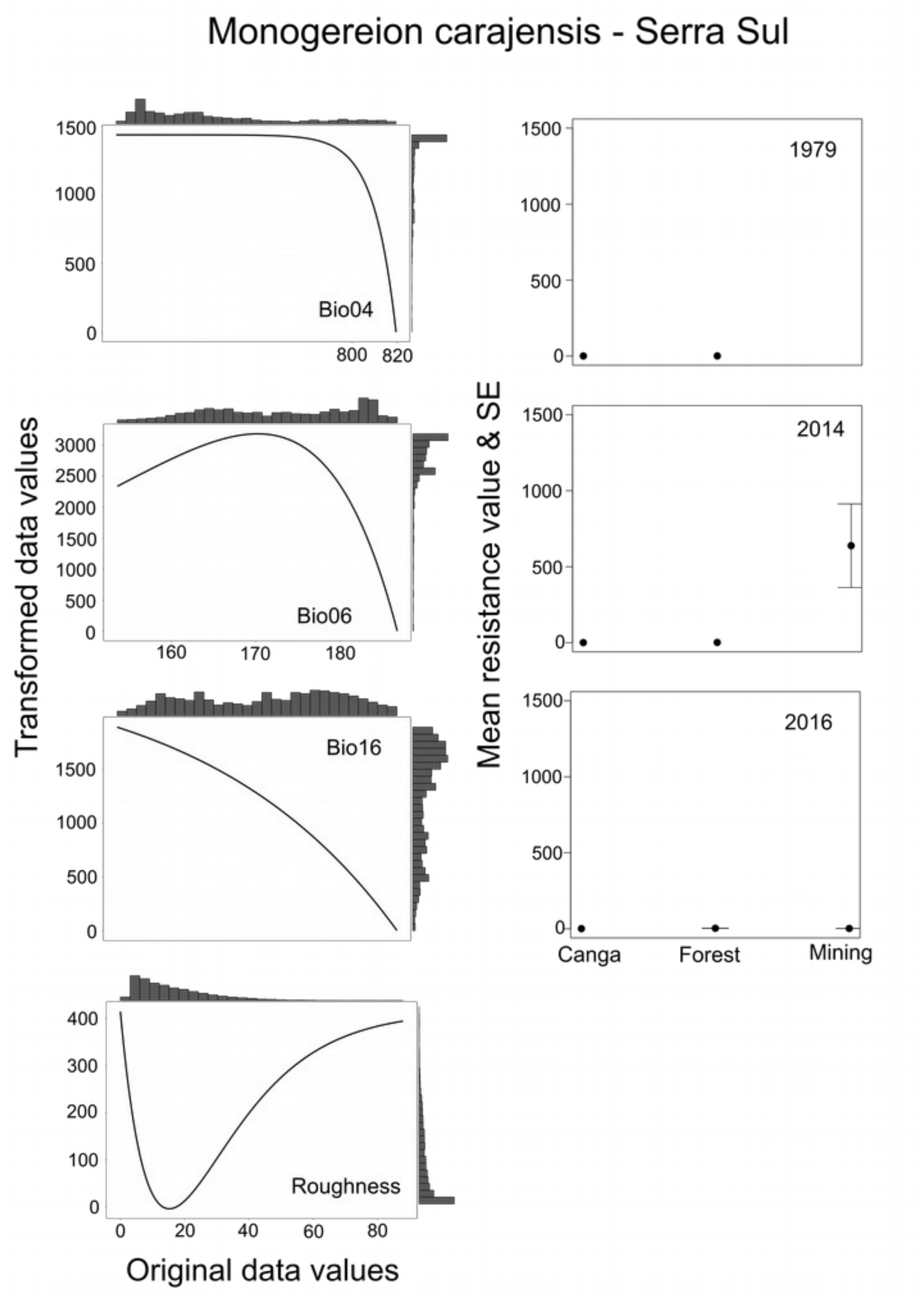


**Figure S10:** Transformation applied to each of continuous (left panels) and categorical (right panels) variables to generate resistance surfaces for *Monogereion carajensis* from Serra Sul. For continuous variables, original data values are represented in x-axes, while transformed data values are shown in y-axes, and histograms present the distribution of original untransformed and transformed variables. For categorical variables, land cover classes were averaged across ten replicates and standard errors (SE) are provided. Categorical resistance surfaces were built for different years (1979, 2014 and 2016). Bio04 and Bio06 represent temperature seasonality and the minimum temperature of the coldest month, respectively, while Bio16 signifies precipitation of wettest quarter.

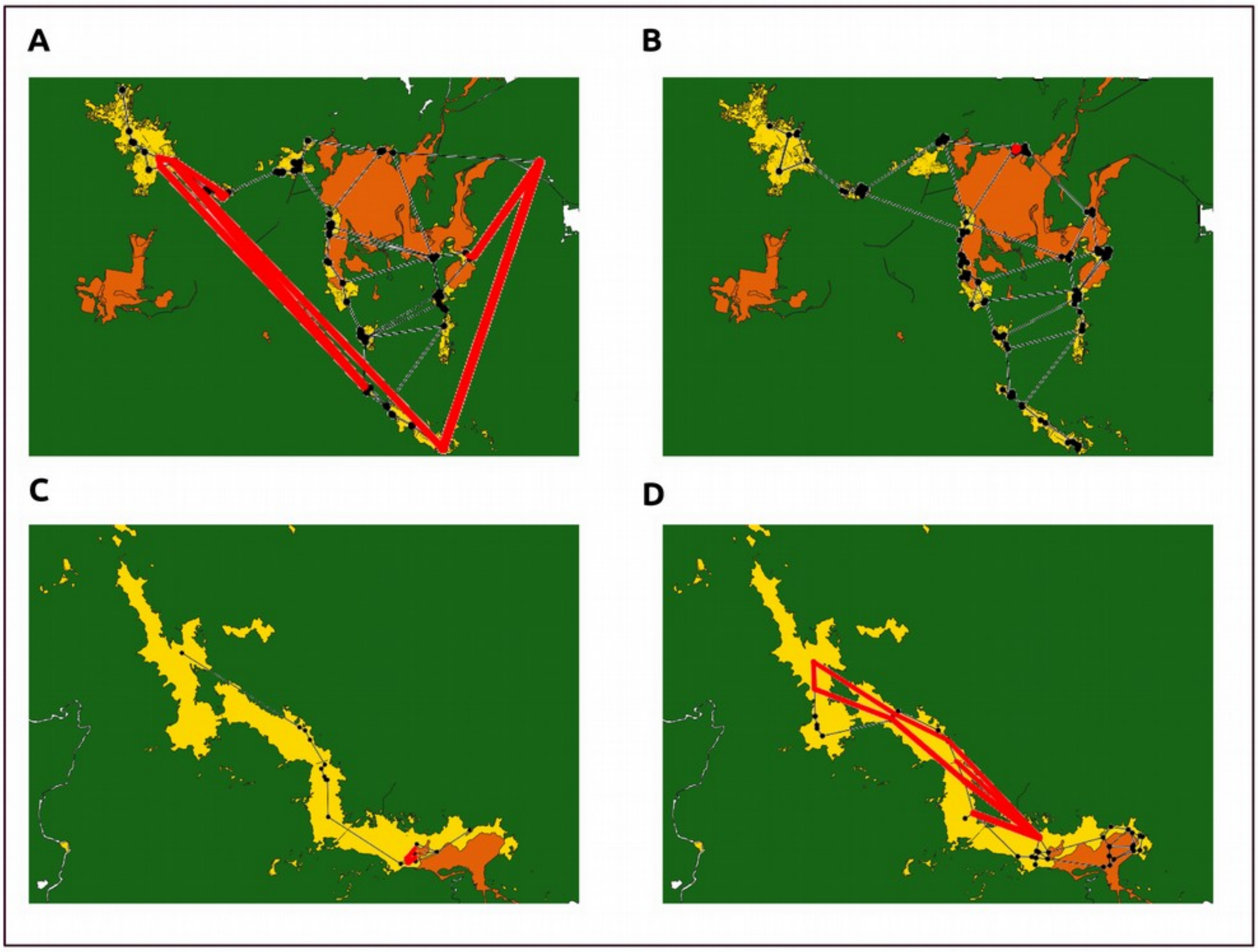


**Figure S11:** Genetic barrier analyses based on Monmonier’s maximum difference algorithm and a Gabriel's graph for *Brasilianthus carajensis* (A and C) and *Monogereion carajensis* (B and D) samples collected in Serra Norte (upper panels) and Serra Sul (lower panels; land cover legends are shown in Fig. 1).

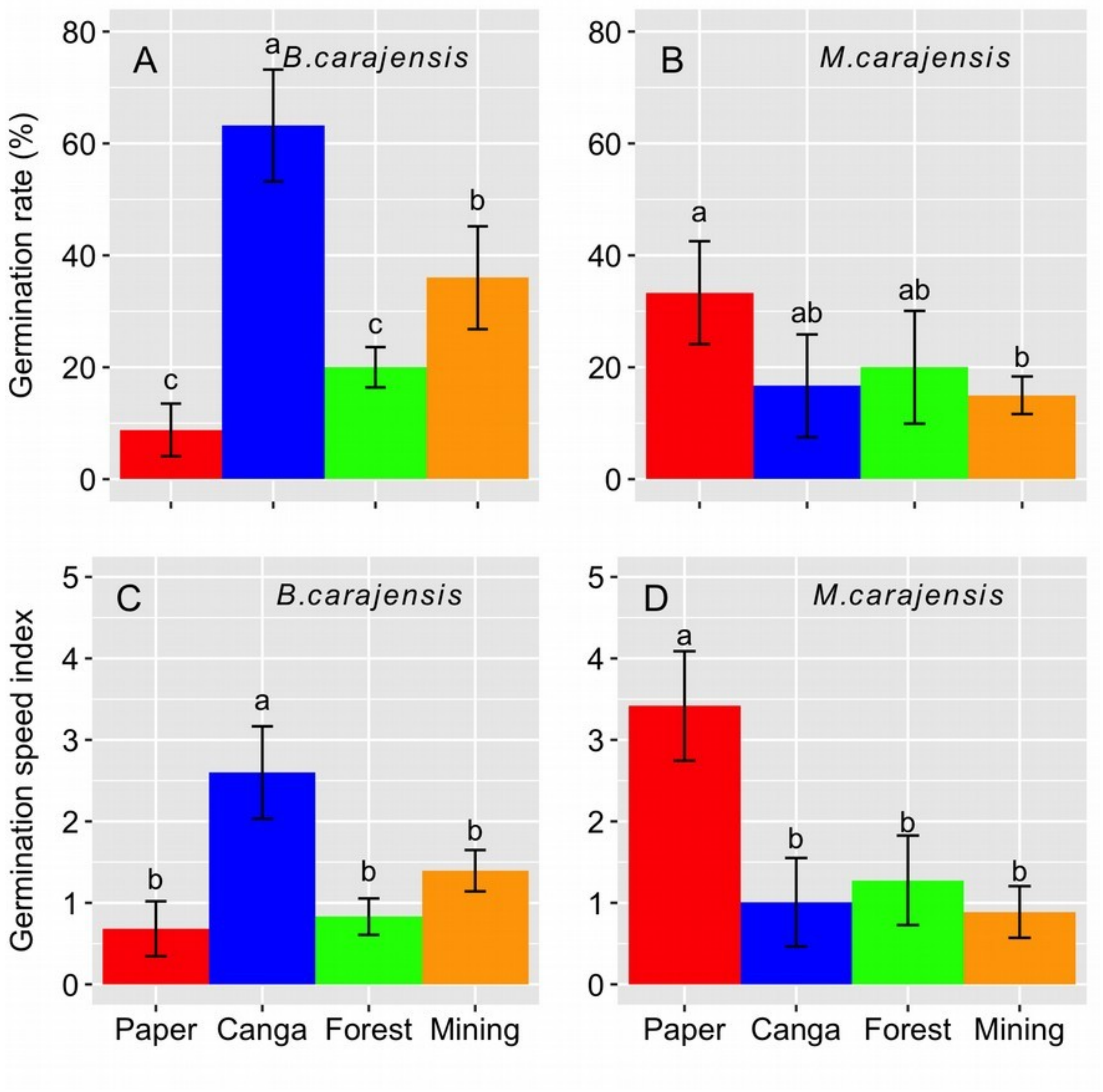


**Figure S12:** Germination rate (A, B) and germination speed (C, D) of *Brasilianthus carajensis* (left panels) and *Monogereion carajensis* (right panels) in four different substrates: Whatman® paper, Canga topsoil, Forest topsoil and Mining waste. Mean values of the cumulative germination rate (percentage of seeds germinated at 33$^{th}$ day) carrying the same letters indicate no significant differences between the treatments after a post hoc Tukey HSD test at $P < 0.05$. The error bars represent the mean standard deviation (n=5).

### Supplementary Tables

**Table S1:** Model selection summary showing the best-fitting genetic diversity models ($\Delta AIC \leq 2$) for *Brasilianthus carajensis* and *Monogereion carajensis* in Serra Norte and Serra Sul, constructed with different buffer sizes. All models contained individual-level genetic diversity metrics (*H* and *f*) as response variable and habitat amount in 2016 as predictors. Buffers sizes varied from 100 m to 900 m, by each 100 m.

| Region | Species | Genetic parameter [b] | Buffer size | AIC | ΔAIC | Weight |
|---|---|---|---|---|---|---|
| Serra Norte | *B. carajensis* [a] | *H* | 100 m | -569.70 | 0.00 | 0.21 |
| | | | 200 m | -568.50 | 1.19 | 0.11 |
| | | | 900 m | -568.50 | 1.21 | 0.11 |
| | | | 800 m | -568.50 | 1.22 | 0.11 |
| | | | 700 m | -568.30 | 1.41 | 0.10 |
| | | | 300 m | -567.90 | 1.75 | 0.08 |
| | | | 600 m | -567.90 | 1.76 | 0.08 |
| | | | 500 m | -567.70 | 1.98 | 0.07 |
| | | | 400 m | -567.70 | 2.00 | 0.07 |
| | | *f* | 900 m | -49.10 | 0.00 | 0.35 |
| | | | 800 m | 48.50 | 0.60 | 0.26 |
| | | | 700 m | 47.60 | 1.52 | 0.16 |
| | *M. carajensis* | *H* | 100 m | -518.40 | 0.00 | 0.15 |
| | | | 900 m | -518.00 | 0.44 | 0.12 |
| | | | 200 m | -518.00 | 0.45 | 0.12 |
| | | | 500 m | -518.00 | 0.47 | 0.12 |
| | | | 800 m | -517.70 | 0.69 | 0.10 |
| | | | 600 m | -517.70 | 0.75 | 0.10 |
| | | | 300 m | -517.50 | 0.92 | 0.09 |

| | | | | | | |
|---|---|---|---|---|---|---|
| | | | 700 m | -517.50 | 0.95 | 0.09 |
| | | | 400 m | -517.40 | 1.08 | 0.09 |
| | | *f* | 400 m | -186.60 | 0.00 | 0.15 |
| | | | 500 m | -186.50 | 0.12 | 0.15 |
| | | | 300 m | -186.40 | 0.21 | 0.14 |
| | | | 600 m | -186.10 | 0.51 | 0.12 |
| | | | 700 m | -185.90 | 0.67 | 0.11 |
| | | | 800 m | -185.90 | 0.75 | 0.10 |
| | | | 900 m | -185.70 | 0.92 | 0.09 |
| | | | 200 m | -185.00 | 1.62 | 0.07 |
| Serra Sul | *B. carajensis* | *H* | 100 m | -74.70 | 0.00 | 0.14 |
| | | | 200 m | -74.60 | 0.07 | 0.13 |
| | | | 300 m | -74.40 | 0.32 | 0.11 |
| | | | 900 m | -74.20 | 0.51 | 0.10 |
| | | | 600 m | -74.10 | 0.57 | 0.10 |
| | | | 800 m | -74.10 | 0.58 | 0.10 |
| | | | 700 m | -74.10 | 0.59 | 0.10 |
| | | | 500 m | -74.10 | 0.61 | 0.10 |
| | | | 400 m | -74.00 | 0.67 | 0.10 |
| | | *f* | 500 m | -5.50 | 0.00 | 0.14 |
| | | | 900 m | -5.50 | 0.00 | 0.14 |
| | | | 800 m | -5.40 | 0.08 | 0.13 |
| | | | 600 m | -5.30 | 0.24 | 0.12 |
| | | | 700 m | -5.20 | 0.30 | 0.12 |
| | | | 400 m | -5.10 | 0.40 | 0.11 |

| | | | | | |
|---|---|---|---|---|---|
| | | 100 m | -4.40 | 1.06 | 0.08 |
| | | 300 m | -4.40 | 1.09 | 0.08 |
| | | 200 m | -4.10 | 1.43 | 0.06 |
| *M. carajensis* | *H* | 900 m | -333.70 | 0.00 | 0.17 |
| | | 800 m | -333.50 | 0.12 | 0.16 |
| | | 700 m | -333.20 | 0.42 | 0.14 |
| | | 600 m | -332.90 | 0.73 | 0.12 |
| | | 500 m | -332.60 | 1.02 | 0.10 |
| | | 400 m | -332.40 | 1.30 | 0.09 |
| | | 300 m | -332.00 | 1.62 | 0.07 |
| | | 200 m | -331.90 | 1.79 | 0.07 |
| | | 100 m | -331.80 | 1.87 | 0.06 |
| | *f* | 900 m | -59.40 | 0.00 | 0.27 |
| | | 800 m | -58.80 | 0.65 | 0.19 |
| | | 700 m | -58.10 | 1.35 | 0.14 |
| | | 600 m | -57.40 | 2.00 | 0.10 |

[a] Genetic cluster was used as random effect to account for the underlying population structure (see methods). [b] Models types are the same as those used in the table S1.

**Table S2:** Model selection summary showing the best-fitting univariate land cover MLPE models (ΔAIC ≤ 2) for *Brasilianthus carajensis* and *Monogereion carajensis* in Serra Norte and Serra Sul using maps with different resolutions (grain size). All models contained inter-individual genetic relatedness as response variable and distances retrieved from the different land cover resistance surfaces as predictors (see methods for details).

| **Region** | **Species** | **Predictor** | **Grain size** | | | | | | | | | | | |
|---|---|---|---|---|---|---|---|---|---|---|---|---|---|---|
| | | | **100 x 100 m** | | | **300 x 300 m** | | | **600 x 600 m** | | | **900 x 900 m** | | |
| | | | **AIC** | **ΔAIC** | **Weight** | **AIC** | **ΔAIC** | **Weight** | **AIC** | **ΔAIC** | **Weight** | **AIC** | **ΔAIC** | **Weight** |
| Serra Norte | *B. carajensis* [a] | Geographic distance | -15453.10 | 0.00 | 1.00 | -15463.20 | 0.00 | 1.00 | -15429.10 | 0.00 | 1.00 | -15469.80 | 0.00 | 1.00 |
| | *M. carajensis* | Land cover 1979 | -23606.80 | 0.00 | 1.00 | -23604.80 | 0.00 | 0.99 | -23594.40 | 0.00 | 1.00 | -23608.00 | 0.00 | 1.00 |
| Serra Sul | *B. carajensis* | Geographic distance | -312.70 | 0.03 | 0.33 | -310.70 | 0.04 | 0.33 | -307.30 | 0.05 | 0.32 | -306.60 | 0.04 | 0.32 |
| | | Land cover 1979 | -312.70 | 0.03 | 0.33 | -310.70 | 0.04 | 0.33 | -307.30 | 0.05 | 0.32 | -306.60 | 0.04 | 0.32 |
| | | Land cover 2014 | -312.80 | 0.00 | 0.33 | -310.80 | 0.00 | 0.33 | -307.40 | 0.00 | 0.33 | -306.70 | 0.00 | 0.32 |
| | *M. carajensis* | Geographic distance | -5681.90 | 0.00 | 0.40 | -5683.30 | 0.00 | 0.40 | -5678.50 | 0.30 | 0.31 | -5676.70 | 0.00 | 0.46 |
| | | Land cover 1979 | -5681.90 | 0.03 | 0.40 | -5683.00 | 0.28 | 0.35 | -5678.80 | 0.00 | 0.36 | -5676.40 | 0.33 | 0.39 |
| | | Land cover 2014 | -5679.90 | 1.94 | 0.15 | -5681.80 | 1.46 | 0.19 | -5677.00 | 1.78 | 0.15 | | | |
| | | Land cover 2016 | | | | | | | -5677.40 | 1.45 | 0.17 | | | |

[a] Genetic cluster was used as random effect to account for the underlying population structure (see methods).

**Table S3:** Model selection summary showing the best-fitting genetic diversity models (ΔAIC ≤ 2) for *Brasilianthus carajensis* and *Monogereion carajensis* in Serra Norte and Serra Sul. All models contained individual-level genetic diversity metrics (*H* and *f*) as response variable and habitat amount in 1979 and habitat loss in 2011, 2014 and 2016 as predictors.

| Region | Species | Response | Predictors [a] | Model type [b] | logLikelihood | AIC | ΔAIC | Weight |
|---|---|---|---|---|---|---|---|---|
| Serra Norte | *B. carajensis* | *H* | None (null model) | LMM [c] | 287.48 | -571.00 | 0.00 | 0.24 |
| | | | Habitat loss in 2016 | | 288.16 | -570.30 | 0.65 | 0.18 |
| | | | Habitat loss in 2014 | | 287.91 | -569.80 | 1.14 | 0.14 |
| | | | Habitat loss in 2011 | | 287.91 | -569.80 | 1.15 | 0.14 |
| | | | Habitat amount in 1979 | | 287.62 | -569.20 | 1.73 | 0.10 |
| | | | Habitat amount in 1979 + Habitat loss in 2016 | | 288.49 | 569.00 | 1.98 | 0.09 |
| | | *f* | Habitat amount in 1979* + Habitat loss in 2016 | LMM [c] | 28.58 | -47.20 | 0.00 | 0.28 |
| | | | Habitat amount in 1979 + Habitat loss in 2011 | | 28.15 | -46.30 | 0.85 | 0.18 |
| | | | Habitat amount in 1979 | | 27.11 | -46.20 | 0.94 | 0.17 |
| | *M. carajensis* | *H* | Habitat amount in 1979* | LMM allowing different variances across Canga plateaus | 410.92 | -799.80 | 0.00 | 0.31 |
| | | | Habitat amount in 1979 + Habitat loss in 2011 | | 411.02 | -798.00 | 1.79 | 0.13 |
| | | | None (null model) | | 408.98 | -798.00 | 1.87 | 0.12 |
| | | | Habitat amount in 1979 + Habitat loss in 2014 | | 410.95 | -797.90 | 1.94 | 0.12 |
| | | | Habitat amount in 1979 + Habitat loss in 2016 | | 410.94 | -797.90 | 1.96 | 0.12 |
| | | *f* | Habitat amount in 1979*** | LMM allowing different variances across Canga plateaus | 126.91 | -231.80 | 0.00 | 0.41 |
| | | | Habitat amount in 1979** + Habitat loss in 2011 | | 127.32 | -230.60 | 1.18 | 0.23 |
| | | | Habitat amount in 1979** + Habitat loss in 2016 | | 127.09 | -230.20 | 1.63 | 0.18 |
| | | | Habitat amount in 1979*** + Habitat loss in 2014 | | 126.93 | -229.90 | 1.95 | 0.15 |
| Serra Sul | *B. carajensis* | *H* | None (null model) | GLS | 40.01 | -76.00 | 0.00 | 0.51 |
| | | | Habitat amount in 1979 | | 40.16 | -74.30 | 1.69 | 0.22 |
| | | *f* | Habitat amount in 1979** | GLS allowing | 13.01 | -14.00 | 0.00 | 0.41 |

| | | | | | | | |
|---|---|---|---|---|---|---|---|
| | | Habitat amount in 1979* + Habitat loss in 2016 | different variances across Canga plateaus | 13.98 | -14.00 | 0.06 | 0.40 |
| | | Habitat loss in 2016* | | 12.03 | -12.10 | 1.96 | 0.15 |
| *M. carajensis* | *H* | Habitat amount in 1979 | GLS allowing different variances across Canga plateaus | 182.62 | -353.20 | 0.00 | 0.39 |
| | | None (null model) | | 180.85 | -351.70 | 1.52 | 0.18 |
| | | Habitat amount in 1979 + Habitat loss in 2014 | | 182.66 | -351.30 | 1.91 | 0.15 |
| | | Habitat amount in 1979 + Habitat loss in 2016 | | 182.62 | -351.20 | 1.99 | 0.14 |
| | *f* | Habitat amount in 1979* | GLS | 32.88 | -59.80 | 0.00 | 0.29 |
| | | Habitat amount in 1979* + Habitat loss in 2014 | | 33.76 | -59.50 | 0.24 | 0.25 |
| | | Habitat amount in 1979* + Habitat loss in 2016 | | 33.67 | -59.30 | 0.43 | 0.23 |

[a] Likelihood Ratio Test (LRT) were performed to assess if each predictor variable significantly improved the model's log-likelihood. Significance levels are highlighted with: *$p < 0.05$; **$p < 0.01$; and ***$p < 0.001$. [b] LMM means linear mixed-effect models and GLS generalized least-squares models. [c] Genetic cluster was used as random effect in this model to account for the underlying population structure (see methods).

**Table S4:** Model-averaged coefficients for the set of best-fitting genetic diversity models (ΔAIC ≤ 2) for *Brasilianthus carajensis* and *Monogereion carajensis* in Serra Norte and Serra Sul. Parameter estimates for each predictor variable are shown followed by standard errors (SE), z-values, p-values and 95% confidence intervals (CI). Cases where CI did not contain zero are highlighted with an asterisk (*).

| Region | Species | Genetic parameter | Predictor | Estimate | SE | *z*-value | *p*-value | CI |
|---|---|---|---|---|---|---|---|---|
| Serra Norte | *B. carajensis* | *H* | Habitat amount in 1979 | $1.00 \times 10^{-3}$ | $4.00 \times 10^{-3}$ | 0.26 | 0.79 | [-0.01,0.01] |
| | | | Habitat loss in 2016 | $-4.00 \times 10^{-3}$ | $9.00 \times 10^{-3}$ | 0.46 | 0.64 | [-0.02,0.01] |
| | | | Habitat loss in 2014 | $-2.00 \times 10^{-3}$ | $8.00 \times 10^{-3}$ | 0.27 | 0.78 | [-0.02,0.01] |
| | | | Habitat loss in 2011 | $-3.99 \times 10^{-3}$ | 0.01 | 0.27 | 0.78 | [-0.02,0.02] |
| | | *f* | Habitat amount in 1979 | 0.20 | 0.09 | 1.99 | 0.04 | [0.01,0.39]* |
| | | | Habitat loss in 2016 | -0.11 | 0.16 | 0.69 | 0.48 | [-0.42,0.20] |
| | | | Habitat loss in 2011 | -0.10 | 0.22 | 0.49 | 0.62 | [-0.54,0.32] |
| | *M. carajensis* | *H* | Habitat amount in 1979 | $4.00 \times 10^{-3}$ | $2.00 \times 10^{-3}$ | 1.49 | 0.13 | [-0.01,0.01] |
| | | | Habitat loss in 2016 | $6.74 \times 10^{-5}$ | $8.57 \times 10^{-4}$ | 0.08 | 0.93 | [-0.01,0.01] |
| | | | Habitat loss in 2014 | $9.08 \times 10^{-5}$ | $9.52 \times 10^{-4}$ | 0.09 | 0.92 | [-0.01,0.01] |
| | | | Habitat loss in 2011 | $1.95 \times 10^{-4}$ | $1.17 \times 10^{-3}$ | 0.17 | 0.86 | [-0.01,0.01] |
| | | *f* | Habitat amount in 1979 | -0.23 | 0.06 | 3.66 | $< 0.001$ | [-0.34,-0.10]* |
| | | | Habitat loss in 2016 | $-7.00 \times 10^{-3}$ | 0.03 | 0.24 | 0.80 | [-0.07,0.05] |
| | | | Habitat loss in 2014 | $-2.00 \times 10^{-3}$ | 0.03 | 0.09 | 0.92 | [-0.05,0.04] |
| | | | Habitat loss in 2011 | -0.01 | 0.04 | 0.35 | 0.72 | [-0.09,0.06] |
| Serra Sul | *B. carajensis* | *H* | Habitat amount in 1979 | $6.00 \times 10^{-3}$ | 0.03 | 0.22 | 0.82 | [-0.04,0.05] |
| | | | Habitat loss in 2016 | $-3.00 \times 10^{-4}$ | 0.01 | 0.02 | 0.98 | [-0.03,0.03] |
| | | *f* | Habitat amount in 1979 | 0.42 | 0.24 | 1.66 | 0.09 | [-0.06,0.89] |
| | | | Habitat loss in 2016 | 0.19 | 0.24 | 0.78 | 0.43 | [-0.27,0.66] |
| | *M. carajensis* | *H* | Habitat amount in 1979 | $-6.00 \times 10^{-3}$ | $5.00 \times 10^{-3}$ | 1.26 | 0.20 | [-0.01,0.01] |
| | | | Habitat loss in 2016 | $-3.03 \times 10^{-5}$ | $1.00 \times 10^{-3}$ | 0.03 | 0.97 | [-0.01,0.01] |
| | | | Habitat loss in 2014 | $5.12 \times 10^{-4}$ | $4.00 \times 10^{-3}$ | 0.12 | 0.90 | [-0.01,0.01] |
| | | *f* | Habitat amount in 1979 | -0.27 | 0.13 | 2.05 | 0.04 | [-0.51,-0.01]* |
| | | | Habitat loss in 2016 | 0.02 | 0.05 | 0.46 | 0.64 | [-0.08,0.13] |
| | | | Habitat loss in 2014 | 0.12 | 0.24 | 0.50 | 0.61 | [-0.34,0.58] |

**Table S5:** Model selection summary showing the best multiple regression MLPE models (ΔAIC ≤ 2) for *Brasilianthus carajensis* and *Monogereion carajensis* in Serra Norte and Serra Sul. All models contained inter-individual genetic relatedness as response variable and the different resistance distances as predictors.

| Region | Species | Predictor [a] | logLikelihood | AIC | ΔAIC | Weight |
|---|---|---|---|---|---|---|
| Serra Norte | *B. carajensis* [b] | Geographic distance*** | 7715.85 | -15421.70 | 0.00 | 0.97 |
| | *M. carajensis* | Land cover 1979*** | 11801.46 | -23594.90 | 0.00 | 0.95 |
| Serra Sul | *B. carajensis* | Roughness*** | 160.67 | -313.30 | 0.00 | 0.31 |
| | | Temperature (Bio6)*** | 160.21 | -312.40 | 0.92 | 0.20 |
| | | Precipitation (Bio19)*** | 160.17 | -312.30 | 0.99 | 0.19 |
| | | Precipitation (Bio16)*** | 159.87 | -311.80 | 1.58 | 0.14 |
| | | Geographic distance*** | 159.80 | -311.60 | 1.73 | 0.13 |
| | *M. carajensis* | Temperature (Bio6)*** | 2843.97 | -5679.90 | 0.00 | 0.40 |
| | | Precipitation (Bio16)*** | 2843.77 | -5679.60 | 0.39 | 0.33 |
| | | Temperature (Bio04)*** | 2843.18 | -5678.40 | 1.58 | 0.18 |

[a] Likelihood Ratio Test (LRT) were performed to assess if each predictor variable significantly improved the model's log-likelihood. Significance levels are highlighted with: *p < 0.05; **p < 0.01; and ***p < 0.001. [b] Genetic cluster was used as random effect to account for the underlying population structure (see methods).

**Table S6:** Coefficients for the set of best-fitting multiple regression MLPE models (ΔAIC ≤ 2) for *Brasilianthus carajensis* and *Monogereion carajensis* in Serra Norte and Serra Sul. Parameter estimates for each predictor variable are shown followed by degrees of freedom (df) standard errors (SE), t-values, p-values and 95% confidence intervals (CI). Cases where CI did not contain zero are highlighted with an asterisk (*).

| Region | Species | Predictor | Estimate | df | SE | *t*-value | *p*-value | CI |
|---|---|---|---|---|---|---|---|---|
| Serra Norte | *B. carajensis* | Geographic distance | -0.02 | 5221 | $8.00 \times 10^{-4}$ | -23.13 | <0.001 | [-0.02,-0.01]* |
| | *M. carajensis* | Land cover 1979 | -0.01 | 7140 | $6.00 \times 10^{-4}$ | -23.57 | < 0.001 | [-0.01,-0.01]* |
| Serra Sul | *B. carajensis* | Roughness | -0.04 | 136 | $7.00 \times 10^{-3}$ | -5.61 | < 0.001 | [-0.05,-0.02]* |
| | | Temperature (Bio6) | -0.04 | 136 | $7.00 \times 10^{-3}$ | -5.51 | < 0.001 | [-0.05,-0.02]* |
| | | Precipitation (Bio19) | -0.04 | 136 | $7.00 \times 10^{-3}$ | -5.49 | < 0.001 | [-0.05,-0.02]* |
| | | Precipitation (Bio16) | -0.04 | 136 | $7.00 \times 10^{-3}$ | -5.46 | < 0.001 | [-0.05,-0.02]* |
| | | Geographic distance | -0.04 | 136 | $7.00 \times 10^{-3}$ | -5.43 | < 0.001 | [-0.05,-0.02]* |
| | *M. carajensis* | Temperature (Bio6) | $-6.1 \times 10^{-3}$ | 1176 | $6.00 \times 10^{-4}$ | -9.76 | < 0.001 | [-0.01,-0.01]* |
| | | Precipitation (Bio16) | $-5.7 \times 10^{-3}$ | 1176 | $5.00 \times 10^{-4}$ | -9.73 | < 0.001 | [-0.01,-0.01]* |
| | | Temperature (Bio04) | $-5.7 \times 10^{-3}$ | 1176 | $5.00 \times 10^{-4}$ | -9.67 | < 0.001 | [-0.01,-0.01]* |
